# A SARS-CoV-2 neutralizing antibody protects from lung pathology in a COVID-19 hamster model

**DOI:** 10.1101/2020.08.15.252320

**Authors:** Jakob Kreye, S Momsen Reincke, Hans-Christian Kornau, Elisa Sánchez-Sendin, Victor Max Corman, Hejun Liu, Meng Yuan, Nicholas C. Wu, Xueyong Zhu, Chang-Chun D. Lee, Jakob Trimpert, Markus Höltje, Kristina Dietert, Laura Stöffler, Niels von Wardenburg, Scott van Hoof, Marie A Homeyer, Julius Hoffmann, Azza Abdelgawad, Achim D Gruber, Luca D Bertzbach, Daria Vladimirova, Lucie Y Li, Paula Charlotte Barthel, Karl Skriner, Andreas C Hocke, Stefan Hippenstiel, Martin Witzenrath, Norbert Suttorp, Florian Kurth, Christiana Franke, Matthias Endres, Dietmar Schmitz, Lara Maria Jeworowski, Anja Richter, Marie Luisa Schmidt, Tatjana Schwarz, Marcel Alexander Müller, Christian Drosten, Daniel Wendisch, Leif E Sander, Nikolaus Osterrieder, Ian A Wilson, Harald Prüss

## Abstract

The emergence of SARS-CoV-2 led to pandemic spread of coronavirus disease 2019 (COVID-19), manifesting with respiratory symptoms and multi-organ dysfunction. Detailed characterization of virus-neutralizing antibodies and target epitopes is needed to understand COVID-19 pathophysiology and guide immunization strategies. Among 598 human monoclonal antibodies (mAbs) from ten COVID-19 patients, we identified 40 strongly neutralizing mAbs. The most potent mAb CV07-209 neutralized authentic SARS-CoV-2 with IC_50_ of 3.1 ng/ml. Crystal structures of two mAbs in complex with the SARS-CoV-2 receptor-binding domain at 2.55 and 2.70 Å revealed a direct block of ACE2 attachment. Interestingly, some of the near-germline SARS-CoV-2 neutralizing mAbs reacted with mammalian self-antigens. Prophylactic and therapeutic application of CV07-209 protected hamsters from SARS-CoV-2 infection, weight loss and lung pathology. Our results show that non-self-reactive virus-neutralizing mAbs elicited during SARS-CoV-2 infection are a promising therapeutic strategy.

## INTRODUCTION

The severe acute respiratory syndrome coronavirus 2 (SARS-CoV-2) started emerging in humans in late 2019, and rapidly spread to a pandemic with millions of cases worldwide. SARS-CoV-2 infections cause coronavirus disease 2019 (COVID-19) with severe respiratory symptoms, but also pathological inflammation and multi-organ dysfunction, including acute respiratory distress syndrome, cardiovascular events, coagulopathies and neurological symptoms (Helms et al., 2020; Zhou et al., 2020; Zhu et al., 2020). Some aspects of the diverse clinical manifestations may result from a hyperinflammatory response, as suggested by reduced mortality in hospitalized COVID-19 patients under dexamethasone therapy (Horby et al., 2020).

Understanding the immune response to SARS-CoV-2 therefore is of utmost importance. Multiple recombinant SARS-CoV-2 mAbs from convalescent patients have been reported (Brouwer et al., 2020; Cao et al., 2020; Ju et al., 2020; Kreer et al., 2020; Robbiani et al., 2020; Rogers et al., 2020; Wec et al., 2020). mAbs targeting the receptor-binding domain (RBD) of the viral spike protein S1 can compete with its binding to human angiotensin converting enzyme 2 (ACE2) and prevent viral entry and subsequent replication (Cao et al., 2020; Ju et al., 2020; Walls et al., 2020). Potent virus neutralizing mAbs that were isolated from diverse variable immunoglobulin (Ig) genes typically carry low levels of somatic hypermutations (SHM). Several of these neutralizing mAbs selected for *in vitro* efficacy showed prophylactic or therapeutic potential in animal models (Cao et al., 2020; Liu et al., 2020; Rogers et al., 2020; Zost et al., 2020). The low number of SHM suggests limited affinity-maturation in germinal centers compatible with an acute infection. Near-germline mAbs usually constitute the first line of defense to pathogens, but carry the risk of self-reactivity to autoantigens (Lerner, 2016; Liao et al., 2011; Zhou et al., 2007). Although critical for the therapeutic use in humans, potential potential tissue-reactivity of near-germline SARS-CoV-2 antibodies has not been examined so far.

Here, we systematically selected 18 strongly neutralizing mAbs out of 598 antibodies from 10 COVID-19 patients by characterization of their biophysical properties, authentic SARS-CoV-2 neutralization, and exclusion of off-target binding to murine tissue. Additionally, we solved two crystal structures of neutralizing mAbs in complex with the RBD, showing antibody engagement with the ACE2 binding site from different approach angles. Finally, we selected mAb CV07-209 by its *in vitro* efficacy and the absence of tissue-reactivity for *in vivo* evaluation. Systemic application of CV07-209 in a hamster model of SARS-CoV-2 infection led to profound reduction of clinical, paraclinical and histopathological COVID-19 pathology, thereby reflecting its potential for translational application in patients with COVID-19.

## RESULTS

### Antibody repertoire analysis of COVID-19 patients

We first characterized the B cell response in COVID-19 using single-cell Ig gene sequencing of human mAbs (Fig. 1A). From ten COVID-19 patients with serum antibodies to the S1 subunit of the SARS-CoV-2 spike protein (Fig. S1, Supplementary Table ST1), we isolated two populations of single cells from peripheral blood mononuclear cells with fluorescence-activated cell sorting (FACS): CD19^+^CD27^+^CD38^+^ antibody secreting cells (ASC) reflecting the unbiased humoral immune response and SARS-CoV-2-S1-labeled CD19^+^CD27^+^ memory B cells (S1-MBC) for characterization of antigen-specific responses (Fig. S2A and S2B). We obtained 598 functional paired heavy and light chain Ig sequences (Supplementary Table ST2). Of 432 recombinantly expressed mAbs, 122 were reactive to SARS-CoV-2-S1 (S1+), with a frequency of 0.0-18.2% (median 7.1%) within ASC and 16.7-84.1% (median 67.1%) within S1-MBC (Fig. 1B and 1C). Binding to S1 did not depend on affinity maturation as measured by the number of SHM (Fig. 1D). Compared to mAbs not reactive to SARS-CoV-2-S1, S1+ mAbs had less SHM, but equal lengths for both their light and heavy chain complementarity-determining region 3 (CDR3) (Fig. S2C, S2D and S2E). Within the ASC and S1-MBC population, 45.0% and 90.2% of S1+ mAbs bound the RBD, respectively (Fig. S2F).

**Fig. 1.**
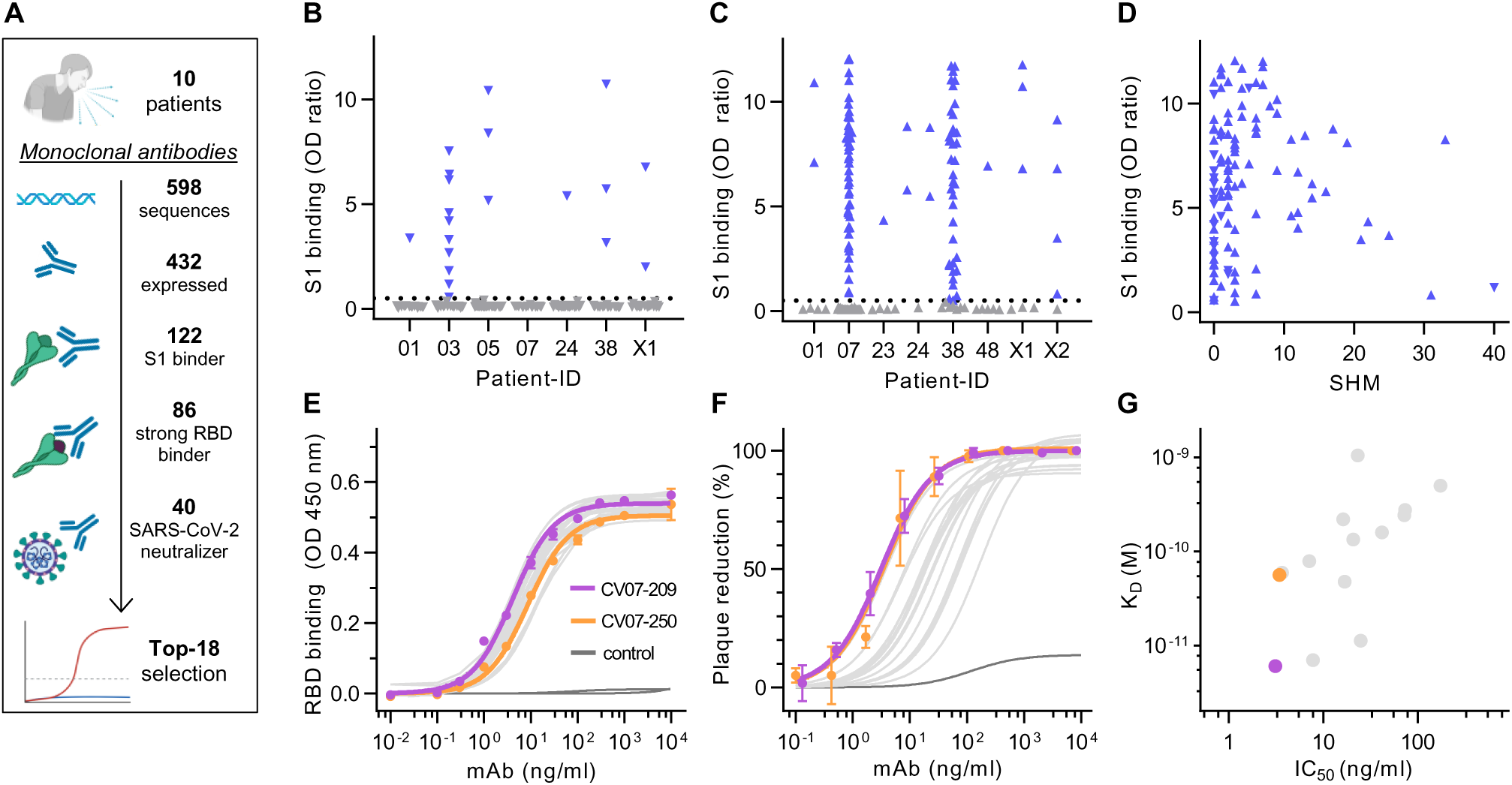
Identification and characterization of potent SARS-CoV-2 neutralizing mAbs. (**A**) Diagram depicting the strategy for isolation of 18 potently neutralizing mAbs (Top-18). (**B**) Normalized binding to S1 of SARS-CoV-2 for mAbs isolated from antibody secreting cells (▾; blue = S1-binding, grey = not S1-binding). (OD=optical density in ELISA) (**C**) Normalized binding to S1 of SARS-CoV-2 for mAbs isolated from S1-stained memory B cells (▴; colors like in (B)) (**D**) S1-binding plotted against the number of somatic hypermutations (SHM) for all S1-reactive mAbs. (**E**) Concentration-dependent binding of Top-18 SARS-CoV-2 mAbs to the RBD of S1 (mean±SD from two wells of one experiment). (**F**) Concentration-dependent neutralization of authentic SARS-CoV-2 plaque formation by Top-18 mAbs (mean±SD from two independent measurements). (**G**) Affinity of mAbs to RBDs (KD determined by surface plasmon resonance) plotted against IC50 of authentic SARS-CoV-2 neutralization.

S1+ mAbs were enriched in certain Ig genes including VH1-2, VH3-53, VH3-66, VK1-33 and VL2-14 (Fig. S3). We identified both clonally related antibody clones within patients and public and shared S1+ clonotypes from multiple patients (Fig. S4). Some public or shared clonotypes had been previously reported, such as IGHV3-53 and IGHV3-66 (Supplementary Table ST3) (Cao et al., 2020; Yuan et al., 2020a), while others were newly identified, such as IGHV3-11 (Fig. S4C).

### Identification and characterization of potent SARS-CoV-2 neutralizing mAbs

We next determined the mAbs with the highest capacity to neutralize SARS-CoV-2 in a plaque reduction neutralization tests (PRNT) using authentic virus (Munich isolate 984) (Wolfel et al., 2020). Of 86 mAbs strongly binding to RBD, 40 showed virus neutralization with a half-maximal inhibitory concentration (IC_50_) ≤250 ng/ml and were considered neutralizing antibodies (Fig. 1A, Supplementary Table ST2), from which 18 (Top-18) were selected for further characterization (Supplementary Table ST4). The antibodies bound to RBD with a half-maximal effective concentration (EC_50_) of 3.8-14.2 ng/ml (Fig. 1E) and an equilibrium dissociation constant (K_D_) of 6.0 pM to 1.1 nM (Fig. S5, Supplementary Table ST4), thereby neutralizing SARS-CoV-2 with an IC_50_ of 3.1-172 ng/ml (Fig. 1F, Supplementary Table ST4). The antibody with the highest affinity, CV07-209, was also the strongest neutralizer (Fig. 1G). We hypothesized that the differences in neutralizing capacity relate to different interactions with the ACE2 binding site. Indeed, the strongest neutralizing mAbs CV07-209 and CV07-250 reduced ACE2 binding to RBD to 12.4% and 58.3%, respectively. Other Top-18 mAbs including CV07-270 interfered only weakly with ACE2 binding (Fig. S6A).

The spike proteins of SARS-CoV-2 and SARS-CoV share more than 70% amino acid sequence identity, whereas sequence identity between SARS-CoV-2 and MERS-CoV and other endemic coronaviruses is significantly lower (Barnes et al., 2020). To analyze potential cross-reactivity of mAbs to other coronaviruses, we tested for binding of the Top-18 mAbs to the RBD of SARS-CoV, MERS-CoV, and the human endemic coronaviruses 229-E, NL63, HKU1 and OC32. CV38-142 detected the RBD of both SARS-CoV-2 and SARS-CoV, whereas no other mAb was cross-reactive to additional coronaviruses (Fig. S7). To further characterize the epitope of neutralizing mAbs, we performed ELISA-based epitope binning experiments using biotinylated antibodies. Co-applications of paired mAbs showed competition of most neutralizing antibodies for RBD binding (Fig. S6B). As an exception, SARS-CoV cross-reactive CV38-142 bound RBD irrespective of the presence of other mAbs, suggesting an independent and conserved target epitope (Fig. S6B).

### Near-germline SARS-CoV-2 neutralizing antibodies can bind to murine tissue

Many SARS-CoV-2 neutralizing mAbs carry few SHM or are in germline-configuration (Fig. 1D) (Ju et al., 2020; Kreer et al., 2020). Such antibodies close to germline might be reactive to more than one target (Zhou et al., 2007). Prompted by the abundance of near-germline SARS-CoV-2 antibodies and to exclude potential side-effects of mAb treatment, we next analyzed whether SARS-CoV-2 antibodies can bind to self-antigens.

Therefore, we tested the binding of S1-mAbs to unfixed murine tissues. Surprisingly, four of the Top-18 potent SARS-CoV-2 neutralizing mAbs showed anatomically distinct tissue reactivities (Fig. 2, Supplementary Table ST4). CV07-200 intensively stained brain sections in the hippocampal formation, olfactory bulb, cerebral cortex and basal ganglia (Fig. 2A). CV07-222 also bound to brain tissue, as well as to smooth muscle (Fig. 2B). CV07-255 and CV07-270 were reactive to smooth muscle from sections of lung, heart, kidney and colon, but not liver (Fig. 2C and 2D, Supplementary Table ST4). None of the Top-18 mAbs bound to HEp-2 cells, cardiolipin or beta-2 microglobulin as established polyreactivity-related antigens (Jardine et al., 2016) (Fig. S8).

**Fig. 2.**
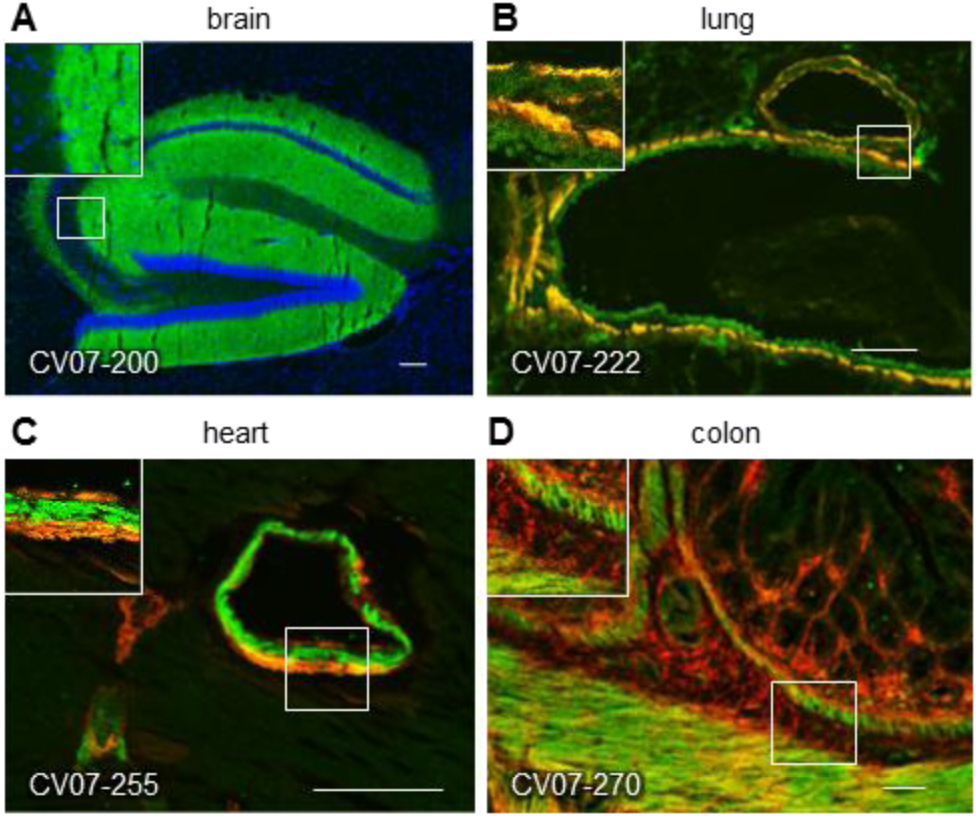
SARS-CoV-2 neutralizing antibodies can bind to murine tissue. Immunofluorescence staining of SARS-CoV-2 mAbs (green) on murine organ sections showed specific binding to distinct anatomical structures, including (**A**) staining of hippocampal neuropil with CV07-200 (cell nuclei depicted in blue), (**B**) staining of bronchial walls with CV07-222, (**C**) staining of vascular walls with CV07-255, and (**D**) staining of intestinal walls with CV07-270. Smooth muscle tissue in (B-D) was co-stained with a commercial smooth muscle actin antibody (red).

### Crystal structures of two mAbs approaching the ACE2 binding site from different angles

For further characterization using X-ray crystallography, we selected two neutralizing mAbs CV07-250 and CV07-270 based on differences in the number of SHM, extent of ACE2 competition and binding to murine tissue. CV07-250 (IC_50_=3.5 ng/ml) had 33 SHM (17/16 on heavy and light chain, respectively), strongly reduced ACE2 binding and showed no binding to murine tissue. In contrast, CV07-270 (IC_50_= 82.3 ng/ml) had only 2 SHM (2/0), did not reduce ACE2 binding in our assay, and showed binding to smooth muscle tissue. Using X-ray crystallography, we determined structures of CV07-250 and CV07-270 in complex with SARS-CoV-2 RBD to resolutions of 2.55 and 2.70 Å, respectively (Fig. 3, Supplementary Tables ST5 and ST6).

**Fig. 3.**
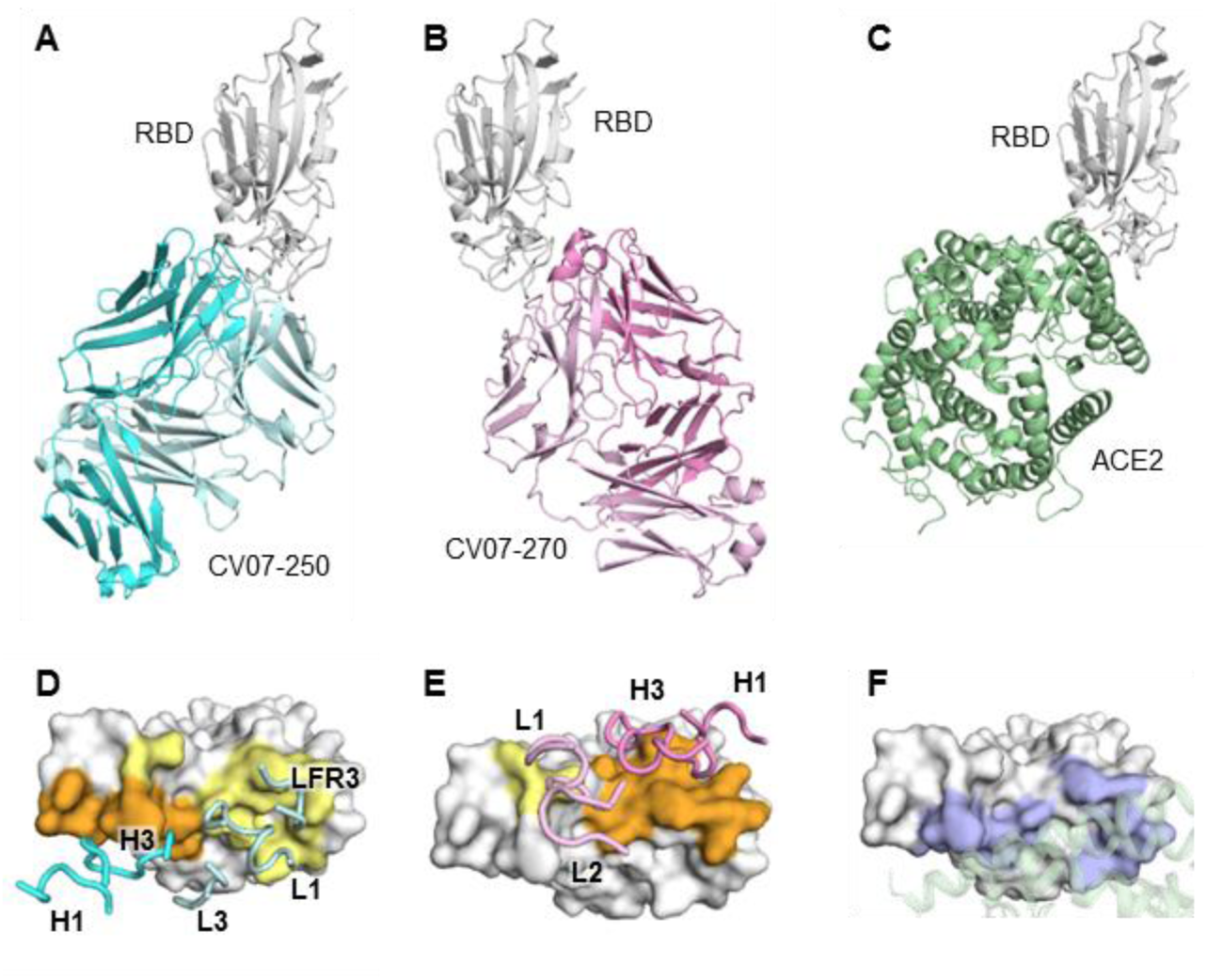
Crystal structures of mAbs in complex with SARS-CoV-2 RBD. (**A**) CV07-250 (cyan) in complex with RBD (white). (**B**) CV07-270 (pink) in complex with RBD (white). (**C**) Human ACE2 with SARS-CoV-2 RBD (PDB 6M0J) (Lan et al., 2020). (**D**-**E**) Epitopes of (**D**) CV07-250 and (**E**) CV07-270. Epitope residues contacting the heavy chain are in orange and the light chain in yellow. CDR loops and framework region that contact the RBD are labeled. (**F**) ACE2-binding residues on the RBD (blue) in the same view as (D) and (E). The ACE2 interacting region is shown in green within a semi-transparent cartoon representation.

The binding mode of CV07-250 to RBD is unusual in that it is dominated by the light chain (Fig. 3A and 3D), whereas in CV07-270, the heavy chain dominates as frequently found in other antibodies (Fig. 3B and 3E). Upon interaction with the RBD, CV07-250 has a buried surface area (BSA) of 399 Å^2^ and 559 Å^2^ on the heavy and light chains, respectively, compared to 714 Å^2^ and 111 Å^2^ in CV07-270. CV07-250 uses CDR H1, H3, L1, L3, and framework region 3 (LFR3) for RBD interaction (Fig. 3D and Fig. 4A, 4B and 4C), whereas CV07-270 interacts with CDR H1, H3, L1, and L2 (Fig. 3E and Fig. 4D, 4E and 4F).

**Fig. 4.**
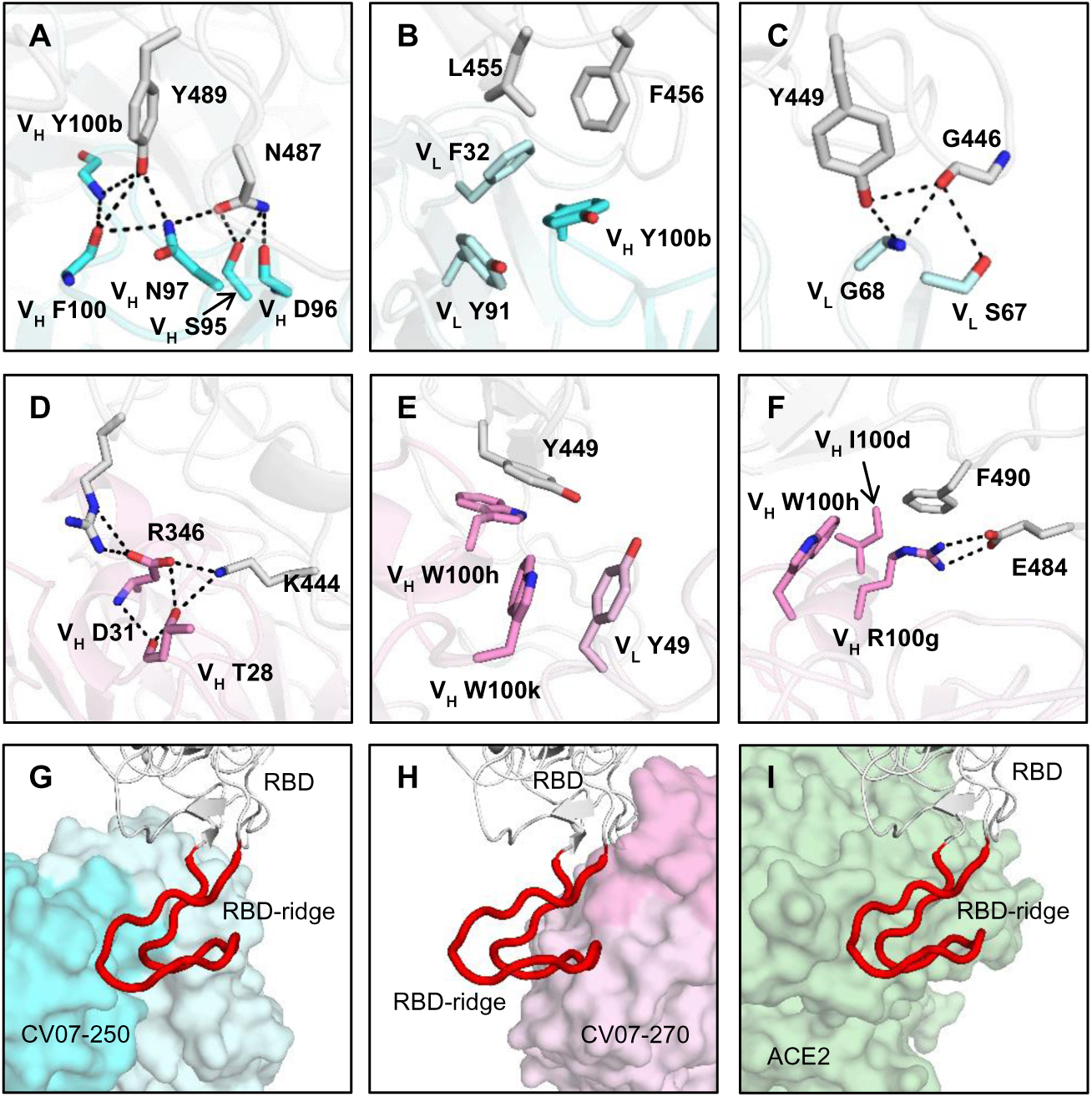
Interactions and angle of approach at the RBD-antibody interface. (**A-C**) Key interactions between CV07-250 (cyan) and RBD (white) are highlighted. (**A**) CDR H3 of CV07-250 forms a hydrogen-bond network with RBD Y489 and N487. (**B**) VH Y100b (CDR H3), VL F32 (CDR L1), and VL Y91 (CDR L3) of CV07-250 form a hydrophobic aromatic patch for interaction with RBD L455 and F456. (**C**) The side chain of VL S67 and backbone amide of VL G68 from FR3 is engaged in a hydrogen-bond network with RBD G446 and Y449. (**D**-**F**) Interactions between CV07-270 (cyan) and RBD (white) are illustrated. (**D**) Residues in CDR H1 of CV07-270 participate in an electrostatic and hydrogen-bond network with RBD R346 and K444. (**E**) VH W100h and VH W100k on CDR H3 of CV07-270 make π-π stacking interactions with Y449. VH W100k is also stabilized by a π-π stacking interaction with VL Y49. (**F**) VH R100g on CDR H3 of CV07-270 forms an electrostatic interaction with RBD E484 as well as a π-cation interaction with RBD F490. Oxygen atoms are in red, and nitrogen atoms in blue. Hydrogen bonds are represented by dashed lines. (**G**-**I**) Zoomed-in views of the different RBD ridge interactions with (**G**) CV07-250, (**H**) CV07-270, and (**I**) ACE2 (PDB 6M0J) (Lan et al., 2020). The ACE2-binding ridge in the RBD is represented by a backbone ribbon trace in red.

The epitope of CV07-250 completely overlaps with the ACE2 binding site with a similar angle of approach as ACE2 (Fig. 3A, 3C, 4G and 4I). In contrast, the CV07-270 epitope only partially overlaps with the ACE2 binding site and the antibody approaches the RBD from a different angle compared to CV07-250 and ACE2 (Fig. 3B, 3C, 4H, 4I), explaining differences in ACE2 competition. Although CV07-250 and CV07-270 both contact 25 epitope residues, only seven residues are shared (G446/G447/E484/G485/Q493/S494/Q498). Furthermore, CV07-270 binds to a similar epitope as SARS-CoV-2 neutralizing antibody P2B-2F6 (Ju et al., 2020) with a similar angle of approach (Fig. S9). In fact, 18 out of 20 residues in the P2B-2F6 epitope overlap with the CV07-270 epitope, although CV07-270 and P2B-2F6 are encoded by different germline genes for both heavy and light chains.

Interestingly, CV07-250 was isolated 19 days after symptom onset, but already acquired 33 SHM, the highest number among all S1+ MBCs (Fig. S2C). Some non-germline amino acids are not directly involved in RBD binding, including all five SHMs on CDR H2 (Fig. S10). This observation suggests that CV07-250 could have been initially affinity-matured against a different antigen.

### Prophylactic and therapeutic mAbs in a COVID-19 animal model

Next, we selected mAb CV07-209 for evaluation of *in vivo* efficacy based on its high capacity to neutralize SARS-CoV-2 and the absence of reactivity to mammalian tissue. We used the hamster model of COVID-19, as it is characterized by rapid weight loss and severe lung pathology (Osterrieder et al., 2020). In this experimental set-up, hamsters were intranasally infected with authentic SARS-CoV-2. Nine hamsters per group received either a prophylactic application of CV07-209 24 hours before viral challenge, or a therapeutic application of CV07-209 or control antibody mGO53 two hours after viral challenge (Fig. 5A).

**Fig. 5.**
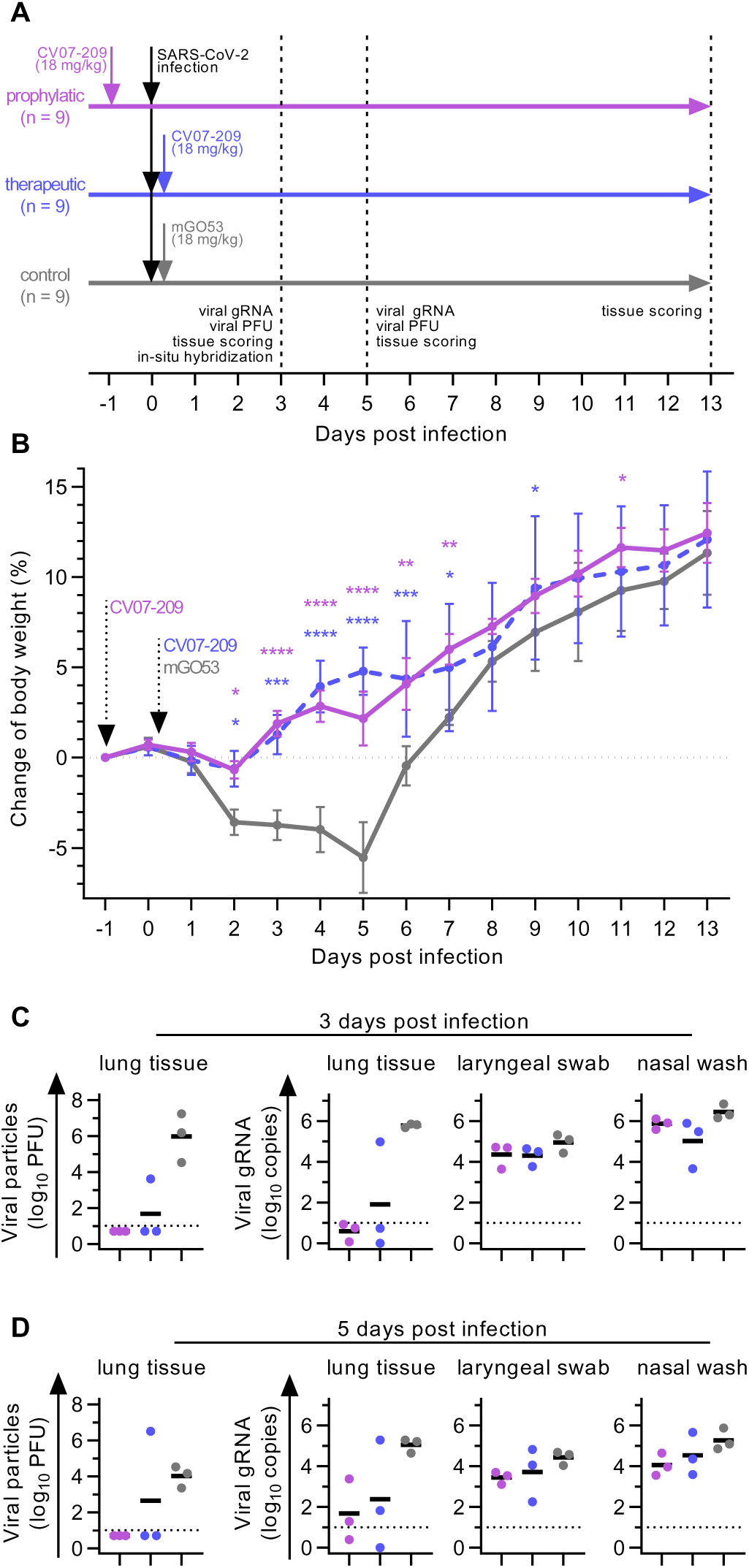
Prophylactic and therapeutic application of mAb CV07-209 in a COVID-19 hamster model. (**A**) Schematic overview of the animal experiment. (**B**) Body weight of hamsters after virus challenge and prophylactic (pink) or therapeutic (blue) application of SARS-CoV-2 neutralizing mAb CV07-209 or control antibody (mean±SEM from n=9 animals per group from day -1 to 3, n=6 from days 4 to 5; n=3 from days 6 to 13; mixed-effects model with posthoc Dunnett’s multiple tests in comparison to control group; significance levels shown as * (p<0.05), ** (p<0.01), *** (p<0.001), **** (p<0.0001), or not shown when not significant). (**C**-**D**) Quantification of plaque forming units (PFU) from lung homogenates and quantification of SARS-CoV-2 RNA copies per 10^5^ cellular transcripts from samples and timepoints as indicated. PFU were set to 5 when not detected, RNA copies below 1 were set to 1. Bars indicate mean. Dotted lines represent detection threshold.

Hamsters under control mAb treatment lost 5.5±4.4% (mean±SD) of body weight, whereas those that received mAb CV07-209 as a therapeutic or prophylactic single dose gained 2.2±3.4% or 4.8±3.4% weight after 5 days post-infection (dpi), respectively. Mean body weights gradually converged in the animals followed up until 13 dpi, reflecting the recovery of control-treated hamsters from SARS-CoV-2 infection (Fig. 5B).

To investigate the presence of SARS-CoV-2 in the lungs, we measured functional SARS-CoV-2 particles from lung tissue homogenates. Plaque forming units were below the detection threshold for all animals in the prophylactic and in 2 of 3 in the treatment group at 3 and 5 dpi (Fig. 5C and 5D). qPCR measurements of lung viral genomic RNA copies revealed a 4-5 and 3-4 log decrease at both time points, indicating a dramatic reduction of SARS-CoV-2 virus particles in both the prophylactic and therapeutic group. However, genomic viral RNA copy numbers from nasal washes and laryngeal swaps were similar between all groups (Fig. 5C and 5D).

Additionally, we performed histopathological analyses of infected hamsters. As expected, all lungs from control-treated animals sacrificed at 3 dpi revealed typical histopathological signs of necro-suppurative pneumonia with suppurative bronchitis, necrosis of bronchial epithelial cells and endothelialitis (Fig. 6A). At 5 dpi, control-treated animals showed marked bronchial hyperplasia, severe interstitial pneumonia with marked type II alveolar epithelial cell hyperplasia and endothelialitis (Fig. 6D). In contrast, animals receiving prophylactic treatment with CV07-209 showed no signs of pneumonia, bronchitis, necrosis of bronchial epithelial cells, or endothelialitis at 3 dpi. A mild interstitial pneumonia with mild type II alveolar epithelial cell hyperplasia became apparent 5 dpi. Animals receiving therapeutic treatment with CV07-209 also showed a marked reduction of histopathological signs of COVID-19 pathology, although at both time points one out of three animals showed mild bronchopulmonary pathology with signs of interstitial pneumonia and endothelialitis. These qualitative findings were mirrored in the reduction of the bronchitis and edema scores (Fig. 6B, 6E and Supplementary Table ST7).

**Fig. 6.**
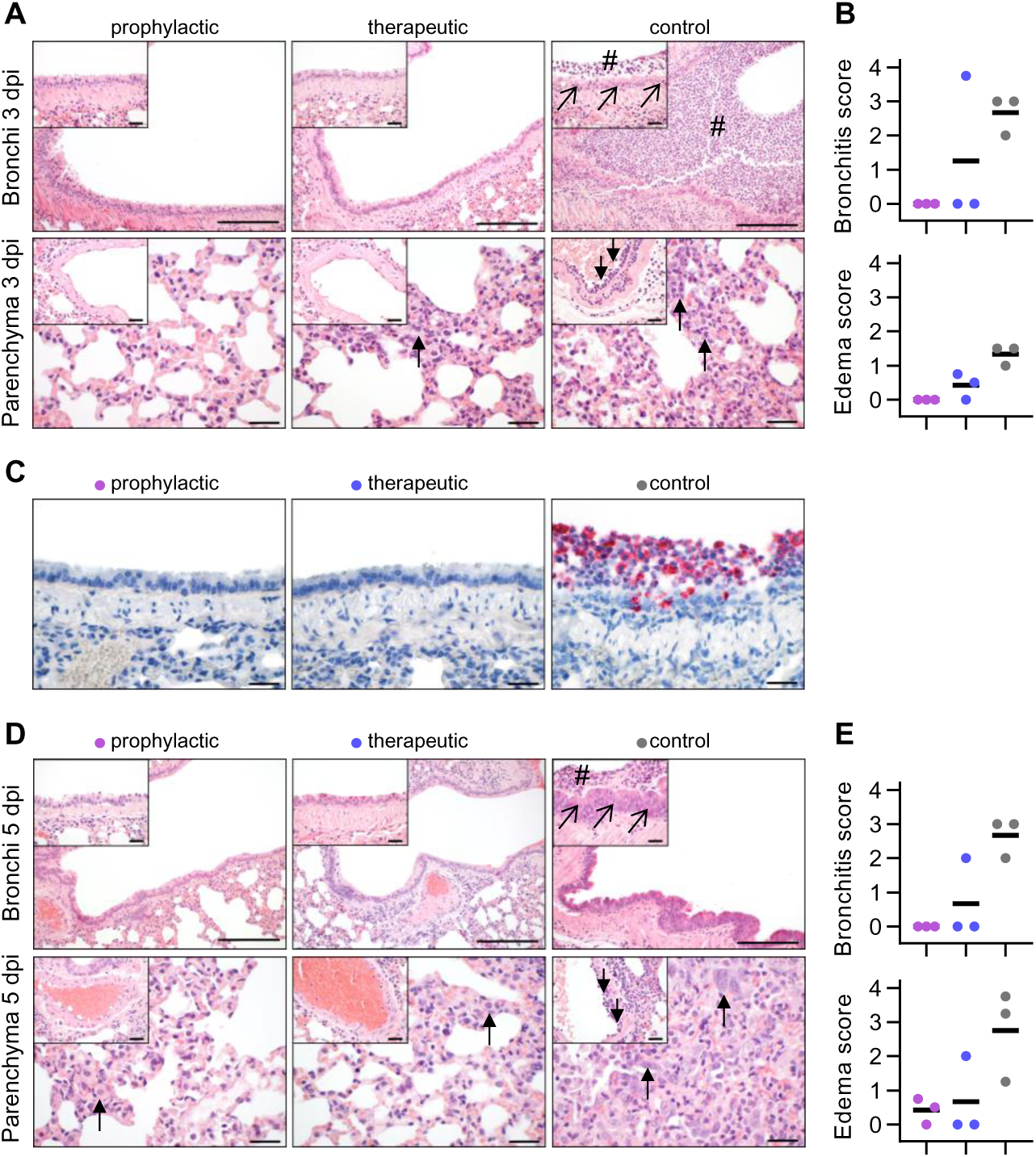
Histopathological analysis of hamsters after SARS-CoV-2 infection. (**A**) Histopathology of representative haematoxylin and eosin stained, paraffin-embedded bronchi with inserted epithelium (upper row) and lung parenchyma with inserted blood vessels (lower row) at 3 dpi. Severe suppurative bronchitis with immune cell infiltration (hash) is apparent only in the control-treated animals with necrosis of bronchial epithelial cells (diagonal arrows). Necro-suppurative interstitial pneumonia (upward arrows) with endothelialitis (downward arrows) is prominent in control-treated animals. Scale bars: 200 µm in bronchus overview, 50 µm in all others. (**B**) Bronchitis and edema score at 3 dpi. (**C**) Detection of viral RNA (red) using *in situ* hybridization of representative bronchial epithelium present only in the control group. Scale bars: 50 µm. (**D**) Histopathology of representative lung sections from comparable areas as in (A) at 5 dpi. Staining of bronchi of control-treated animals showed a marked bronchial hyperplasia with hyperplasia of epithelial cells (diagonal arrow) and still existing bronchitis (hash), absent in all prophylactically treated and in 2/3 therapeutically treated animals (upper row). Lung parenchyma staining of control-treated animals showed severe interstitial pneumonia with marked type II alveolar epithelial cell hyperplasia and endothelialitis (insets, downward arrows). Compared to control-treated animals, prophylactically treated animals showed only mild signs of interstitial pneumonia with mild type II alveolar epithelial cell hyperplasia (upward arrow), whereas therapeutically treated animals showed a more heterogeneous picture with 1/3 showing no signs of lung pathology, 1/3 animal showing only mild signs of interstitial pneumonia, and 1/3 animal showing a moderate multifocal interstitial pneumonia. Scale bars: 200 µm in bronchus overview, 50 µm in all others. (**E**) Bronchitis and edema score at 5 dpi.

To confirm the absence of viral particles under CV07-209 treatment, we performed *in-situ* hybridization of viral RNA at 3 dpi. No viral RNA was detectable in the prophylactic group, whereas all animals in the control group and one in the therapeutic group revealed intensive staining of viral RNA in proximity of bronchial epithelial cells (Fig. 6C). Taken together, these findings show that systemic application of SARS-CoV-2 neutralizing mAb CV07-209 protects hamsters from COVID-19 lung pathology and weight loss in both the prophylactic and the therapeutic setting.

## DISCUSSION

Our results add several new aspects to the growing knowledge on the humoral immune response in SARS-CoV-2 infections. First, we solved the structures of two neutralizing mAbs binding to the RBD of SARS-CoV-2 at resolutions of 2.55 and 2.70 Å, allowing detailed characterization of the target epitopes and gaining insight into the mechanism of mAbs neutralizing SARS-CoV-2. SARS-CoV-2 mAbs can compete with ACE2 binding and exert neutralizing activity by inhibiting viral particle binding to host cells (Barnes et al., 2020; Brouwer et al., 2020; Cao et al., 2020; Ju et al., 2020; Kreer et al., 2020; Robbiani et al., 2020; Rogers et al., 2020; Wec et al., 2020), a key mechanism previously identified in SARS-CoV neutralizing antibodies (Prabakaran et al., 2006; ter Meulen et al., 2006). Steric hindrance of mAbs blocking ACE2 binding to the RBD provides one mechanistic explanation of virus neutralization (Barnes et al., 2020; Cao et al., 2020; Wu et al., 2020). CV07-250 clearly belongs to this category of antibodies, as its epitope lies within the ACE2 binding site and it approaches the RBD from a similar angle as ACE2. In contrast, the epitope of CV07-270 only partially overlaps with the ACE2 binding site and approaches the RBD ridge from a different angle. In line with these findings, competition of CV07-270 with ACE2 binding as detected by ELISA was very weak. Its mechanism of virus neutralization therefore remains elusive. Of note, there have been reports of neutralizing antibodies targeting epitopes distant to the ACE2 binding site (Chi et al., 2020). Future research will need to clarify if additional mechanisms like triggering conformational changes in the spike protein upon antibody binding contribute to virus neutralization, as reported for SARS-CoV (Walls et al., 2019).

Secondly, the majority of our SARS-CoV-2 mAbs are close to germline configuration, supporting previous studies (Kreer et al., 2020; Robbiani et al., 2020). Given the increased probability of auto-reactivity of near-germline antibodies, we examined reactivity with murine tissue and indeed found that a fraction of SARS-CoV-2 neutralizing antibodies also bound to with brain, lung, heart, kidney or gut expressed epitopes. Such reactivity with host antigens should ideally be prevented by immunological tolerance mechanisms, but complete exclusion of such antibodies would generate “holes” in the antibody repertoire. In fact, HIV utilizes epitopes shared by its envelope and mammalian self-antigens, thus harnessing immunological tolerance to impair anti-HIV antibody responses (Yang et al., 2013) and impeding successful vaccination (Jardine et al., 2016). To defy viral escape in HIV, but similarly COVID-19, anergic strongly self-reactive B cells likely enter germinal centers and undergo clonal redemption to mutate away from self-reactivity, while retaining HIV or SARS-CoV-2 binding (Reed et al., 2016). Interestingly, longitudinal analysis of mAbs in COVID-19 showed that the number of SHM in SARS-CoV-2-neutralizing antibodies only marginally increased over time (Kreer et al., 2020). This finding suggests that the self-reactivity observed in this study may not be limited to mAbs of the early humoral immune response in SARS-CoV-2 infections. Whether self-reactive antibodies could contribute to extra-pulmonary symptoms in COVID-19, awaits further studies and should be closely monitored also in vaccination trials.

Finally, we evaluated in detail the *in vivo* efficacy of the most potent neutralizing antibody CV07-209 in a Syrian hamster model. This model is characterized by a severe phenotype including weight loss and distinct lung pathology. Our results demonstrated that prophylaxis and treatment with a single dose of CV07-209 not only led to clinical improvement as shown by the absence of weight loss, but also to markedly reduced lung pathology. While the findings confirm the efficacy of prophylactic mAb administration reported by other groups in mice, hamsters and rhesus macaques (Cao et al., 2020; Liu et al., 2020; Rogers et al., 2020; Zost et al., 2020), our work is the first to demonstrate the efficacy of post-exposure treatment in hamsters leading to viral clearance, clinical remission and prevention of lung injury. To our knowledge, data on post-exposure treatment so far were restricted to transgenic hACE2 mice (Cao et al., 2020) and a mouse model using adenovector delivery of human ACE2 before viral challenge (Liu et al., 2020). Collectively, our results indicate that mAb treatment can be fine-tuned for exclusion of self-reactivity with mammalian tissues and that mAb administration can also be efficacious after the infection, which will be the prevailing setting in COVID-19 patients.

## Acknowledgments

We thank Stefanie Bandura, Matthias Sillmann and Doreen Brandl for excellent technical assistance, Christian Meisel for performing a cardiolipin ELISA and Martin Barner for assistance in generating the circos plot in fig. S4B. We acknowledge BIAFFIN GmbH & Co KG (Kassel, Germany) for performance of SPR measurements and Dr. Désirée Kunkel from Flow & Mass Cytometry Core Facility at Charité - Universitätsmedizin Berlin for support with single cell sorting.

## Author Contributions

JK, SMR, HCK, VMC, JT, KD, HL, MY, NCW, IAW, HP designed the study. SMR, LS, JH, SH, MW, NS, FK, CF, DW and LES collected patient material and led the clinical study PA-COVID-19. JK, SMR, HCK, ESS, VMC, HL, MY, NCW, XZ, CCDL, JT, MH, KD, LS, NvW, SvH, MAH, JH, AA, LDB, DV, LYL, PCB, AH, KS, LMJ, AR, MLS, TS and DW carried out experiments. JK, SMR, HCK, ESS, VMC, HL, MY, NCW, JT, MH, LDB, KD, ADG, ME, DS, ADG, MAM, CD, NO, IAW, HP analyzed and interpreted acquired data. JK, SMR and HP drafted the manuscript. All authors critically revised and approved the submitted version.

## Materials availability

Antibodies will be made available by the corresponding authors upon request under a Material Transfer Agreement (MTA) for non-commercial usage.

## Data and Code Availability

X-ray coordinates and structure factors are deposited at the RCSB Protein Data Bank under accession codes 6XKQ and 6XKP. The raw sequencing data associated with this manuscript will be deposited together with the analysis using BASE on Code Ocean upon publication and will be shared upon request during review. The software used for Ig sequence analysis is available on https://github.com/automatedSequencing/BASE.

## MATERIALS AND METHODS

### SARS-CoV-2-infected individuals and sample collection

The patients have given written informed consent and analyses were approved by the Institutional Review Board of Charité - Universitätsmedizin Berlin, corporate member of Freie Universität Berlin, Humboldt-Universität Berlin, and Berlin Institute of Health, Berlin. All patients in this study were tested positive for SARS-CoV-2 infection by RT-PCR. Patient characteristics are described in Supplementary Table ST1.

### PBMC collection and FACS staining

Recombinant SARS-CoV-2-S1 protein produced in HEK cells (Creative Diagnostics, DAGC091) was covalently labeled using CruzFluor647 (Santa Cruz Biotechnology, sc-362620) according to the manufacturer’s instructions.

Using fluorescence-activated cell sorting we sorted viable single cells from freshly isolated peripheral blood mononuclear cells (PBMCs as 7AAD^-^CD19^+^CD27^+^CD38^+^ antibody-secreting cells (ASCs) or SARS-CoV2-S1-enriched 7AAD^-^CD19^+^CD27^+^ memory B cells (MBCs) into 96-well PCR plates. Staining was performed on ice for 25 minutes in PBS with 2 % FCS using the following antibodies: 7- AAD 1:400 (Thermo Fisher Scientific), CD19-BV786 1:20 (clone SJ25C1, BD Biosciences, 563326), CD27-PE 1:5 (clone M-T271, BD Biosciences, 555441), CD38-FITC 1:5 (clone HIT2, BD Biosciences, 560982), and SARS-CoV-S1-CF647 at 1 µg/ml for patients CV07, CV38, CV23, CV24, CV 38, CV48, CV-X1, CV-X2 and CV01 (second time point, Fig. S1). The first patients (CV01 (first time point), CV03, and CV05) were stained with a divergent set of antibodies: CD19-PE 1:50 (clone HIB19, BioLegend, 302207), CD38-PEcy7 1:50 (clone HIT2, BioLegend, 303505), CD27-APC 1:50 (clone O323, BioLegend, 302809) and DAPI as viability dye.

### Generation of recombinant human monoclonal antibodies

Monoclonal antibodies were generated following our established protocols (Kornau et al., 2020; Kreye et al., 2016) with modifications as mentioned. We used a nested PCR strategy to amplify variable domains of immunoglobulin heavy and light chain genes from single cell cDNA and analyzed sequences with aBASE module of customized Brain Antibody Sequence Evaluation (BASE) software (Reincke et al., 2019). Pairs of functional Ig genes were PCR-amplified using specific primers with Q5 Polymerase (NEB). PCR-product and linearized vector were assembled using Gibson cloning with HiFi DNA Assembly Master Mix (NEB). Cloning was considered successful when sequence identify >99.5% was given, verified by the cBASE module of BASE software. For mAb expression, human embryonic kidney cells (HEK293T) were transiently transfected with matching Ig heavy and light chains. Three days later mAb containing cell culture supernatant was harvested. Ig concentrations were determined and used for reactivity and neutralization screening, if Ig concentration was above 1 µg/ml. For biophysical characterization assays and *in vivo* experiments, supernatants were purified using Protein G Sepharose beads (GE Healthcare), dialyzed against PBS and sterile-filtered using 0.2 µm filter units (GE Healthcare). For *in vivo* experiments, mAbs were concentrated using Pierce™ 3K Protein Concentrator PES (Thermo Scientific).

### SARS-CoV-2-S1 ELISA

Screening for SARS-CoV-2-specific mAbs was done by using anti-SARS-CoV-2-S1 IgG ELISAs (EUROIMMUN Medizinische Labordiagnostika AG) according to the manufacturer’s protocol. mAb containing cell culture supernatants were pre-diluted 1:5, patient sera 1:100. Optical density (OD) ratios were calculated by dividing the OD at 450 nm by the OD of the calibrator included in the kit. OD ratios of 0.5 were considered reactive.

### RBD ELISA

Binding to the receptor-binding domain (RBD) of S1 was tested in an ELISA. To this end, a fusion protein (RBD-Fc) of the signal peptide of the NMDA receptor subunit GluN1, the RBD-SD1 part of SARS-CoV2-S1 (amino acids 319-591) and the constant region of rabbit IgG1 heavy chain (Fc) was expressed in HEK293T cells and immobilized onto 96-well plates from cell culture supernatant via anti-rabbit IgG (Dianova, 111-035-045) antibodies. Then, human mAbs were applied and detected using horseradish peroxidase (HRP)-conjugated anti-human IgG (Dianova, 109-035-088) and the HRP substrate 1-step Ultra TMB (Thermo Fisher Scientific, Waltham, MA). All S1+ mAbs were screened at a human IgG concentration of 10 ng/ml to detect strong RBD binders and the ones negative at this concentration were re-evaluated for RBD reactivity using a 1:5 dilution of the cell culture supernatants. To test for specificity within the coronavirus family, we expressed and immobilized Fc fusion proteins of the RBD-SD1 regions of MERS-CoV, SARS-CoV and the endemic human coronaviruses HCoV-229-E, HCoV-NL63, HCoV-HKU1, and HCoV-229E and applied mAbs at 1 µg/ml. The presence of immobilized antigens was confirmed by incubation with HRP-conjugated anti-rabbit IgG (Dianova). Assays for concentration-dependent RBD binding (Fig. 1E) were developed using 1-step Slow TMB (Thermo Fisher Scientific). EC_50_ was determined from non-linear regression models using Graph Pad Prism 8.

To evaluate the ability of mAbs to interfere with the binding of ACE2 to SARS-CoV-2 RBD, we expressed ACE2-HA, a fusion protein of the extracellular region of human ACE2 (amino acids 1-615) followed by a His-tag and a hemagglutinin (HA)-tag in HEK293T cells and applied it in a modified RBD-ELISA. Captured RBD-Fc was incubated with mAbs at 0.5 µg/ml for 15 minutes and subsequently with ACE2-HA-containing cell culture supernatant for 1 h. ACE2-HA binding was detected using anti-HA antibody HA.11 (clone 16B12, BioLegend, San Diego, CA, MMS-101P), HRP-conjugated anti-mouse IgG (Dianova, 115-035-146) and 1-step UltraTMB.

For experiments regarding the competition between mAbs for RBD binding, purified monoclonal antibodies were biotinylated using EZ-Link Sulfo-NHS-Biotin (Thermo Fisher) according to the manufacturer’s instructions. Briefly, 50-200 µg of purified antibody were incubated with 200-fold molar excess Sulfo-NHS-Biotin for 30 minutes at room temperature. Excess Sulfo-NHS-Biotin was removed by dialysis for 16 hours. Recovery rate of IgG ranged from 60-100%. RBD-Fc captured on ELISA plates was incubated with mAbs at 10 µg/ml for 15 minutes. Then, one volume of biotinylated mcAbs at 100 ng/ml was added and the mixture incubated for additional 15 minutes, followed by detection using HRP-conjugated streptavidin (Roche Diagnostics) and 1-step Ultra TMB. Background by the HRP-conjugated detection antibodies alone was subtracted from all absorbance values.

### Circos plot of public clonotypes

Antibodies which share same V and J gene on both Ig heavy and light chain are considered to be one clonotype. Such clonotypes are considered *public* if they are identified in different patients. After identification of public clonotypes, they were plotted in a Circos plot using the R package circlize (Gu et al., 2014).

### Identification of 18 strongly neutralizing antibodies

To identify the most potent SARS-CoV-2 neutralizing mAb, all 122 S1-reactive mAbs were screened for binding to RBD. 86 were defined as strongly binding to RBD (defined as detectable binding at 10 ng/ml in an RBD ELISA) and then assessed for neutralization of authentic SARS-CoV-2 at 25 and 250 ng/ml using mAb-containing cell culture supernatants. Antibodies were further selected (i) as the strongest neutralizing mAb of the respective donor and / or (ii) with an estimated IC_50_ of 25 ng/ml or below and / or (iii) with an estimated IC_90_ of 250 ng/ml or below. These were defined as the 18 most potent antibodies (Top-18) and expressed as purified antibodies for detailed biophysical characterization.

### Surface plasmon resonance measurements

The antigen (SARS-CoV-2 S protein-RBD-mFc, Accrobiosystems) was reversibly immobilized on a C1 sensor chip via anti-mouse IgG. Purified mAbs were injected at different concentrations in a buffer consisting of 10 mM HEPES pH 7.4, 150 mM NaCl, 3 mM EDTA, 0.05% Tween 20. CV-X1-126 and CV38-139 were analyzed in a buffer containing 400 mM NaCl as there was a slight upward drift at the beginning of the dissociation phase due to non-specific binding of to the reference flow. Multi-cycle-kinetics analyses were performed in duplicates except for non-neutralizing CV03-191. K_a_, K_d_ and K_D_- values were determined using a monovalent analyte model. Recordings were performed on a Biacore T200 instrument at 25°C.

### Plaque reduction neutralization test

To detect neutralizing activity of SARS-CoV-2-specific mAbs, plaque reduction neutralization tests (PRNT) were done as described before (Wolfel et al., 2020). Briefly, Vero E6 cells (1.6 x10^5^ cells/well) were seeded in 24-well plates and incubated overnight. For each dilution step, mAbs were diluted in OptiPro and mixed 1:1 with 200 μl virus (Munich isolate 984) (Wolfel et al., 2020) solution containing 100 plaque forming units. The 400 μl mAb-virus solution was vortexed gently and incubated at 37°C for 1 hour. Each 24-well was incubated with 200 μl mAb-virus solution. After 1 hour at 37°, the supernatants were discarded and cells were washed once with PBS and supplemented with 1.2% Avicel solution in DMEM. After 3 days at 37°C, the supernatants were removed and the 24-well plates were fixed and inactivated using a 6% formaldehyde/PBS solution and stained with crystal violet. All dilutions were tested in duplicates. For PRNT-screening mAb dilutions of 25 and 250 ng of IgG/ml were assessed. IC_50_ was determined from non-linear regression models using Graph Pad Prism 8.

### Immunocytochemistry

Recombinant spike protein-based immunofluorescence assays were done as previously described (Buchholz et al., 2013; Corman et al., 2020; Wolfel et al., 2020). Briefly, VeroB4 cells were transfected with previously described pCG1 plasmids encoding SARS-CoV-2, MERS-CoV, HCoV- NL63, -229E, -OC43, and -HKU1 spike proteins (Buchholz et al., 2013). For transfection, Fugene HD (Roche) was used in a Fugene to DNA ratio of 3:1. After 24 hours, transfected as well as untransfected VeroB4 cells were harvested and resuspended in DMEM/10% FCS to achieve a cell density of 2.5×10^5^ cells/ml each. Transfected and untransfected VeroB4 cells were mixed 1:1 and 50 μl of the cell suspension was applied to each incubation field of a multitest cover slide (Dunn Labortechnik). The multitest cover slides were incubated for 6 hours before they were washed with PBS and fixed with ice-cold acetone/methanol (ratio 1:1) for 10 minutes. For the immunofluorescence test, the incubation fields were blocked with 5% non-fat dry milk in PBS/0.2% Tween for 60 minutes. Purified mAbs were diluted in EUROIMMUN sample buffer to a concentration of 5 µg/ml and 30 μl of the dilution was applied per incubation field. After 1 hour at room temperature, cover slides were washed 3 times for 5 minutes with PBS/0.2% Tween. Secondary detection was done using a 1:200 dilution of a goat-anti human IgG-Alexa488 (Dianova). After 30 minutes at room temperature, slides were washed 3 times for 5 minutes and rinsed with water. Slides were mounted using DAPI prolonged mounting medium (FisherScientific).

### Crystal structure determination of Fab-RBD complexes

The receptor binding domain (RBD; residues 319-541) of the SARS-CoV-2 spike (S) protein was expressed in High Five cells and purified using affinity and size exclusion chromatography as described previously (Yuan et al., 2020b). Residues in the elbow region of CV07-250 (112SSASTKG118) were mutated to 112FNQIKP117 to reduce elbow flexibility and facilitate crystal packing (Bailey et al., 2018). CV07-250 and CV07-270 Fabs were expressed in ExpiCHO cells and purified using affinity and size exclusion chromatography. The Fab/RBD complexes were formed by mixing the two components in an equimolar ratio and incubating overnight at 4°C before setting-up crystal trials. The complexes of CV07-250/RBD and CV07-270/RBD were screened for crystallization at 20.0 and 12.0 mg/ml, respectively, using 384 conditions of the JCSG Core Suite (Qiagen) on our robotic CrystalMation system (Rigaku) at The Scripps Research Institute. Crystals appeared after day 3, were harvested after day 7, and then flash-cooled in liquid nitrogen for X-ray diffraction experiments. Diffraction data were collected at cryogenic temperature (100 K) at Stanford Synchrotron Radiation Lightsource (SSRL) on the newly constructed Scripps/Stanford beamline 12-1 with a beam wavelength of 0.97946 Å and processed with HKL3000 (Minor et al., 2006). Diffraction data were collected from crystals grown in conditions: 0.085 M HEPES pH 7.5, 10% (v/v) ethylene glycol, 15% (v/v) glycerol, 8.5% (v/v) 2-propanol, 17% (w/v) polyethylene glycol 4000 for the CV07-250/RBD complex and 0.1 M sodium cacodylate pH 6.5, 0.2 M sodium chloride, 2 M ammonium sulfate, 15% (v/v) ethylene glycol for the CV07-270/RBD complex. The X-ray structures were solved by molecular replacement (MR) using PHASER (McCoy et al., 2007) with MR models for the RBD and CV07-270 Fab from PDB 6W41 (Yuan et al., 2020b) and 4FQH, respectively. The MR model for CV07-250 Fab was generated using Repertoire Builder (Schritt et al., 2019). Iterative model building and refinement were carried out in COOT (Emsley and Cowtan, 2004) and PHENIX (Adams et al., 2010), respectively. In the CV07-250 + RBD structure, residues 319-337, 357- 366, 371-374, 383- 396, 516-541 were not modeled due to paucity of electron density. The N and C terminal regions are normally disordered in the SARS CoV-2 RBD structures. These flexible regions are not involved in any other contacts, including crystal lattice contacts, and are on the opposite side of the RBD to the epitope, which is well ordered. In the CV07-270 + RBD structure, Fab residues in a region of the heavy-chain constant domain also have greater mobility as commonly found in Fabs. Likewise, the N and C-terminal residues of 319-333 and 528-541 in both RBD molecules of the asymmetric unit are disordered. Epitope and paratope residues, as well as their interactions, were identified by accessing PDBePISA at the European Bioinformatics Institute (https://www.ebi.ac.uk/pdbe/prot_int/pistart.html).

### Murine tissue reactivity screening

Preparations of brain, lung, heart, liver, kidney and gut from 8-12 weeks old C57BL/6J mice were frozen in -50°C 2-methylbutane, cut on a cryostat in 20 µm sections and mounted on glass slides. For tissue reactivity screening according to established protocols (Kreye 2016 Brain), thawed unfixed tissue slices were rinsed with PBS then blocked with blocking solution (PBS supplemented with 2% Bovine Serum Albumin (Roth) and 5% Normal Goat Serum (Abcam)) for 1 hour at room temperature before incubation of mAbs at 5 µg/ml overnight at 4°C. After three PBS washing steps, goat anti-human IgG-Alexa Fluor 488 (Dianova, 109-545-003) diluted in blocking solution was applied for 2 hours at room temperature before additional three washes and mounting using DAPI-containing Fluoroshield. Staining was examined under an inverted fluorescence microscope (Olympus CKX41, Leica DMI6000) or confocal device (Leica TCS SL). For co-staining, tissue was processed as above, but sections were fixed with 4% PFA in PBS for 10 minutes at room temperature before blocking. For co-staining, the following antibodies were used: mouse Smooth Muscle Actin (clone 1A4, Agilent, 172 003), goat anti-mouse IgG-Alexa Fluor 594 (Dianova, 115-585-003). For nuclei staining DRAQ5™ (abcam, ab108410) was used.

### HEp2 cell assay

HEp-2 cell reactivity was investigated using the NOVA Lite HEp-2 ANA Kit (Inova Diagnostics) according to the manufacturer’s instructions using mAb containing culture supernatant (screening of all S1+ mAbs) or purified mAbs at 50 µg/ml (polyreactivity testing of CV07-200, CV07-209, CV07-222, CV07-255, CV07-270 and CV38-148) and examined under an inverted fluorescence microscope.

### Polyreactivity screening ELISA

Purified mAbs were screened for reactivity against cardiolipin and beta-2 microglobulin at 50 µg/ml using routine laboratory ELISAs kindly performed by Christian Meisel (Labor Berlin).

### Hamster model of SARS-CoV-2 infection

The animal experiment was approved by the Landesamt für Gesundheit und Soziales in Berlin, Germany (approval number 0086/20) and performed in compliance with relevant national and international guidelines for care and humane use of animals. In vitro and animal work was conducted under appropriate biosafety precautions in a BSL-3 facility at the Institute of Virology, Freie Universität Berlin, Germany. Twenty-seven six-week old female and male golden Syrian hamsters (Mesocricetus auratus; outbred hamster strain RjHan:AURA, Janvier Labs) were kept in groups of 3 animals in enriched, individually ventilated cages. The animals had ad libitum access to food and water and were allowed to acclimate to these conditions for seven days prior to prophylactic treatment and infection. Cage temperatures and relative humidity were recorded daily and ranged from 22-24°C and 40-55%, respectively.

Virus stocks for animal experiments were prepared from the previously published SARS-CoV-2 München isolate (Wolfel et al., 2020). Viruses were propagated on Vero E6 cells (ATCC CRL-1586) in minimal essential medium (MEM; PAN Biotech) supplemented with 10% fetal bovine serum (PAN Biotech) 100 IU/ml Penicillin G and 100 µg/ml Streptomycin (Carl Roth). Stocks were stored at -80°C prior to experimental infections.

For the SARS-CoV-2 challenge experiments, hamsters were randomly distributed into three groups: In the first group (prophylaxis group), animals received an intraperitoneal (i.p.) injection of 18 mg per kg bodyweight of SARS-CoV-2 neutralizing mAb CV07-209 24 hours prior to infection. In the second and third group (treatment and control group, respectively), animals were given the identical mAb amount two hours after infection, either with 18 mg/kg of CV07-209 (treatment group) or with 20 mg/kg of non-reactive isotype-matched mGO53 (control group). Hamsters were infected intranasally with SARS-CoV-2 infected as previously described (Osterrieder et al., 2020).

On days 2, 5 and 13 post infection, three hamsters of each group were euthanized by exsanguination under general anesthesia as described (Nakamura et al., 2017). Nasal washes, tracheal swabs, and lungs (left and right) were collected for histopathological examinations and/or virus titrations and RT-qPCR. Body weights were recorded daily and clinical signs of all animals were monitored twice daily throughout the experiment.

### Histopathology and in situ hybridization

For histopathological examination and *in situ* hybridization (ISH) of lung tissues, the left lung lobe was carefully removed and immersed in fixative solution (4% formalin, pH 7.0) for 48 hours. Lungs were then embedded in paraffin and cut in 2 μm sections. For histopathology, slides were stained with hematoxylin and eosin (HE) followed by blinded microscopic evaluation by board-certified veterinary pathologists as previously described (Dietert et al., 2017; Osterrieder et al., 2020). The following parameters were evaluated to assess three different scores: (1) the bronchitis score that includes severity of bronchial inflammation and epithelial cell necrosis of bronchi; (2) the regeneration score combining the presence of hyperplasia of bronchial epithelial cells and alveolar type II epithelial cells; and (3) the edema score including alveolar and perivascular edema.

ISH was performed as reported previously (Erickson et al., 2020; Osterrieder et al., 2020) using the ViewRNA™ ISH Tissue Assay Kit (Invitrogen by Thermo Fisher Scientific) following the manufacturer’s instructions with minor adjustments. Probes for the detection of the Nucleoprotein (N) gene RNA of SARS-CoV-2 (NCBI database NC_045512.2, nucleotides 28,274 to 9,533, assay ID: VPNKRHM), and the murine housekeeping gene eukaryotic translation elongation factor-1α (EF1a; assay ID: VB1-14428-VT, Affymetrix, Inc.), that shares 95% sequence identity with the Syrian hamster, were designed. Lung sections (2 µm thickness) on adhesive glass slides were dewaxed in xylol and dehydrated in ethanol. Tissues were incubated at 95°C for 10 minutes with subsequent protease digestion for 20 minutes. Sections were fixed with 4% paraformaldehyde in PBS (Alfa Aesar, Thermo Fisher) and hybridized with the probes. Amplifier and label probe hybridizations were performed according to the manufacturer’s instructions using fast red as the chromogen, followed by counterstaining with hematoxylin for 45 s, washing in tap water for 5 minutes, and mounting with Roti®-Mount Fluor-Care DAPI (4, 6-diaminidino-2-phenylindole; Carl Roth). An irrelevant probe for the detection of pneumolysin was used as a control for sequence-specific binding. HE-stained and ISH slides were analyzed and images taken using a BX41 microscope (Olympus) with a DP80 Microscope Digital Camera and the cellSens™ Imaging Software, Version 1.18 (Olympus).

### Virus titrations, RNA extractions and RT-qPCR

To determine virus titers from 25 mg lung tissue, tissue homogenates were serially diluted and titrated on Vero E6 cells in 12-well-plates. Three days later, cells were formalin-fixed, stained with crystal violet and plaques were counted. RNA was extracted from homogenized lungs, nasal washes and tracheal swabs using the innuPrep Virus DNA/RNA Kit (Analytik Jena) according to the manufacturer’s instructions. We quantified RNA using a one-step RT qPCR reaction with the NEB Luna Universal Probe One-Step RT-qPCR kit (New England Biolabs) and previously published TaqMan primers and probes (SARS-CoV-2 E_Sarbeco and hamster RPL18) (Corman et al., 2020; Zivcec et al., 2011) on a StepOnePlus RealTime PCR System (Thermo Fisher Scientific).

### Statistical Analysis

All statistical tests were performed using GraphPad Prism, version 8.4. For comparison of SHM number, ordinary one-way ANOVA tests with posthoc Tukey’s multiple comparisons tests were used. Statistical significance of bodyweight changes from hamster experiments was tested using a mixed-effects model (two-way ANOVA) with posthoc Dunnett’s multiple comparisons test in comparison to control group.

## SUPPLEMENTARY MATERIALS

**Fig. S1.**
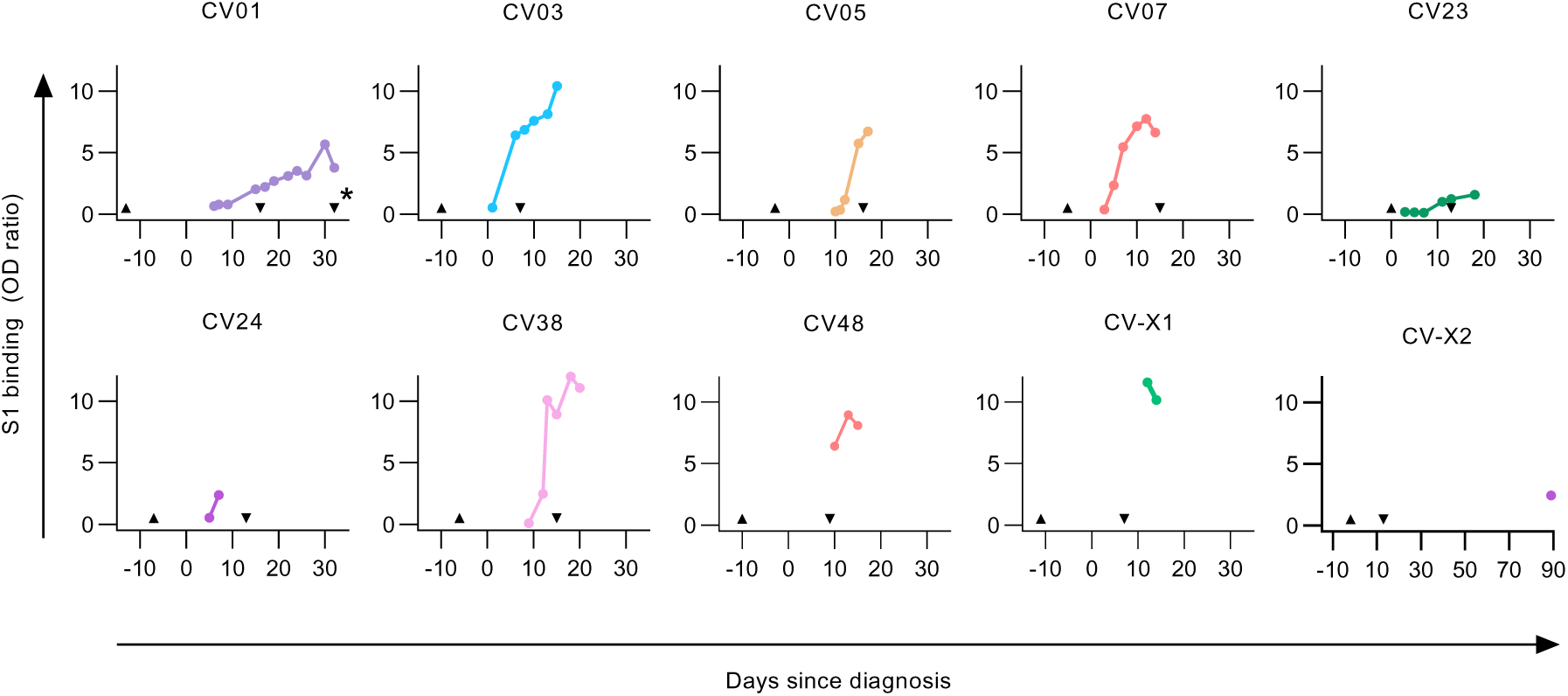
SARS-CoV-2-S1 serum IgG response from COVID-19 patients. Serum IgG response determined as the normalized optical density (OD) in a SARS-CoV-2-S1 ELISA in relation to the time point of diagnosis defined by the first positive qPCR test. Upward arrowhead denotes the appearance of first symptoms. Downward arrowhead denotes the PBMC isolation. From patient CV01, PBMC samples were isolated at two time points as indicted by the second downward arrow with an asterisk (*).

**Fig. S2.**
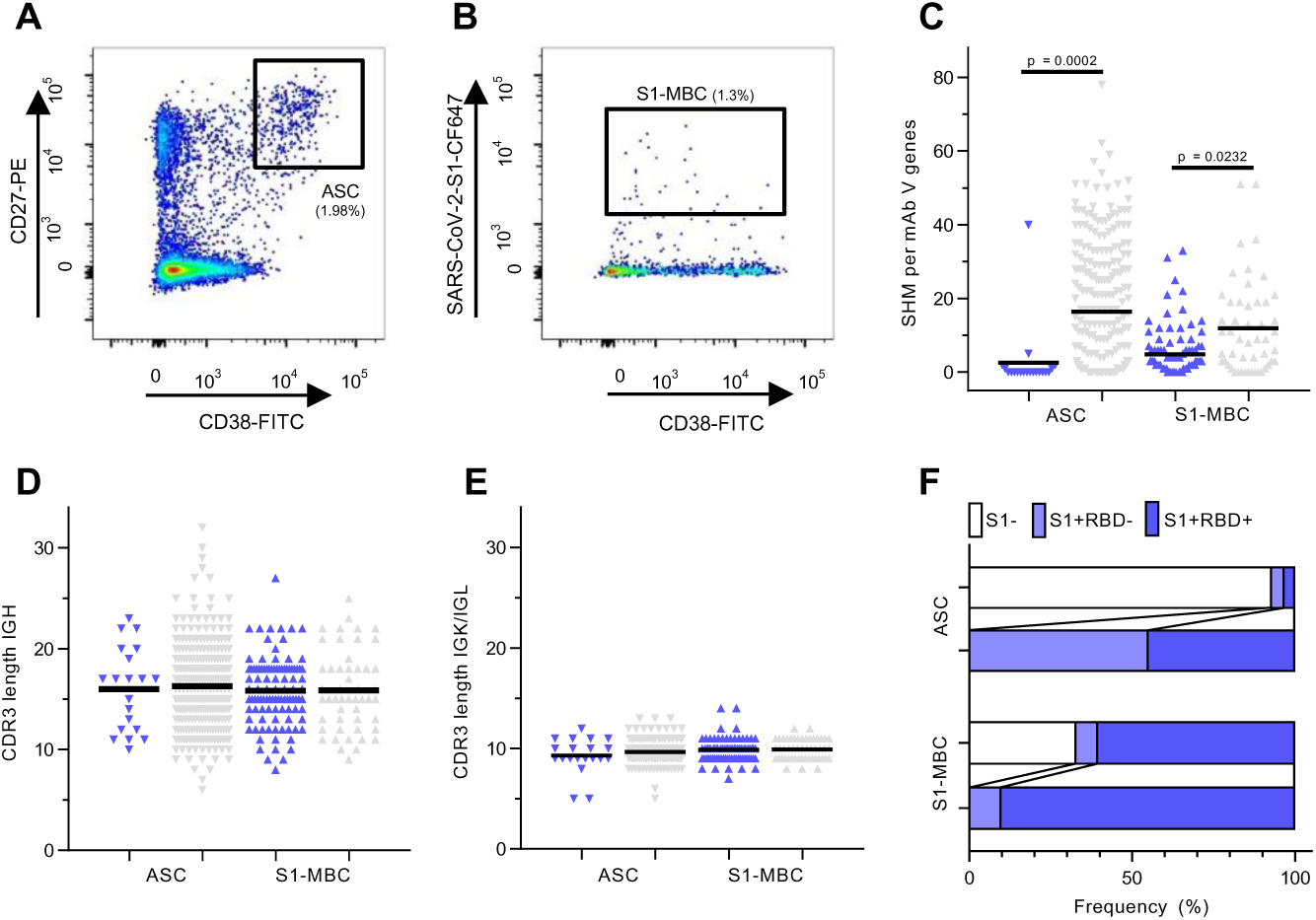
Flow cytometry gating and characteristics of immunoglobulin sequences. (**A**-**B**) A representative flow cytometry plot from patient CV38 indicating gating on (**A**) CD19^+^CD27^+^antibody-secreting cells (ASC) and (**B**) SARS-CoV-2-S1-stained memory B cells (S1-MBC). Cells were pre-gated on live CD19^+^ B cells. (**C**) Comparison of somatic hypermutation (SHM) count within immunoglobulin V genes combined from heavy and light chains of S1-reactive (S1+, blue) and non-S1-reactive (S1-, grey) mAbs. Statistical significance was determined using a one-way ANOVA (F = 19.22) with posthoc Tukey’s multiple comparisons tests. (ASC: n = 20 S1+, n = 260 S1-; S1-MBC: n = 102 S1+, n = 50 S1-). All expressed mAbs are displayed. Each triangle represents a S1+ (blue) or S1- (grey) mAb, isolated from an ASC (pointing downwards) or a S1-MBC (pointing upwards). (**D**-**E**) Length comparison of complementarity-determining region (CDR) 3 amino acid sequences between S1+ and S1- mAbs within (**D**) heavy and (**E**) light chains. Symbols and colors have the same meaning as in (C). (**F**) Frequency of RBD-binder (S1+RBD+) and non-RBD-binder (S1+RBD-) relative to all expressed mAbs (upper lanes) and relative to S1+ mAbs (lower lanes).

**Fig. S3.**
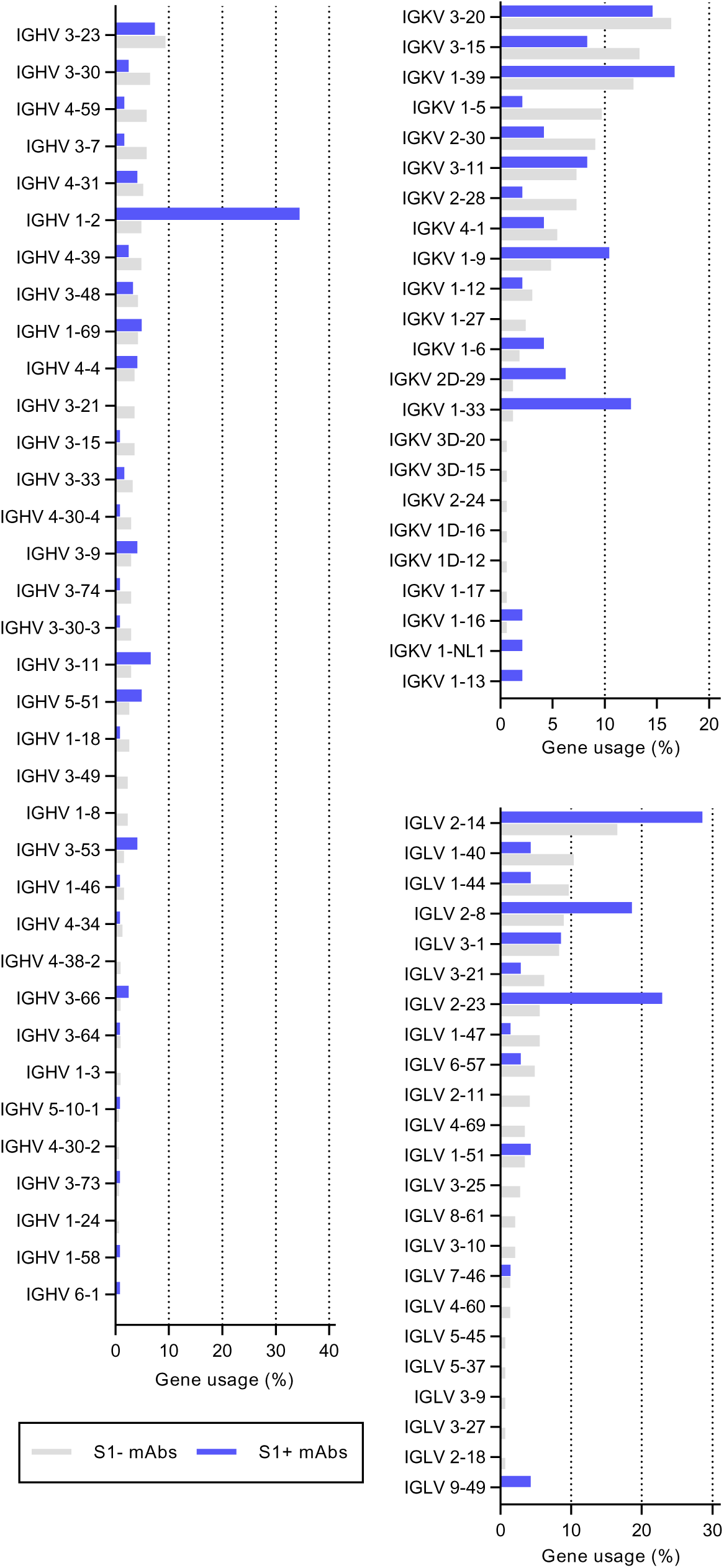
Comparison of variable gene usage. Comparison of gene usage between SARS-CoV-2-S1-reactive (S1+) and non-reactive (S1-) mAbs is shown for immunoglobulin (**A**) variable heavy (IGHV), (**B**) variable kappa (IGKV) and (**C**) variable lambda (IGLV) genes. Bars depict percentage of gene usage of all expressed mAbs within each group.

**Fig. S4.**
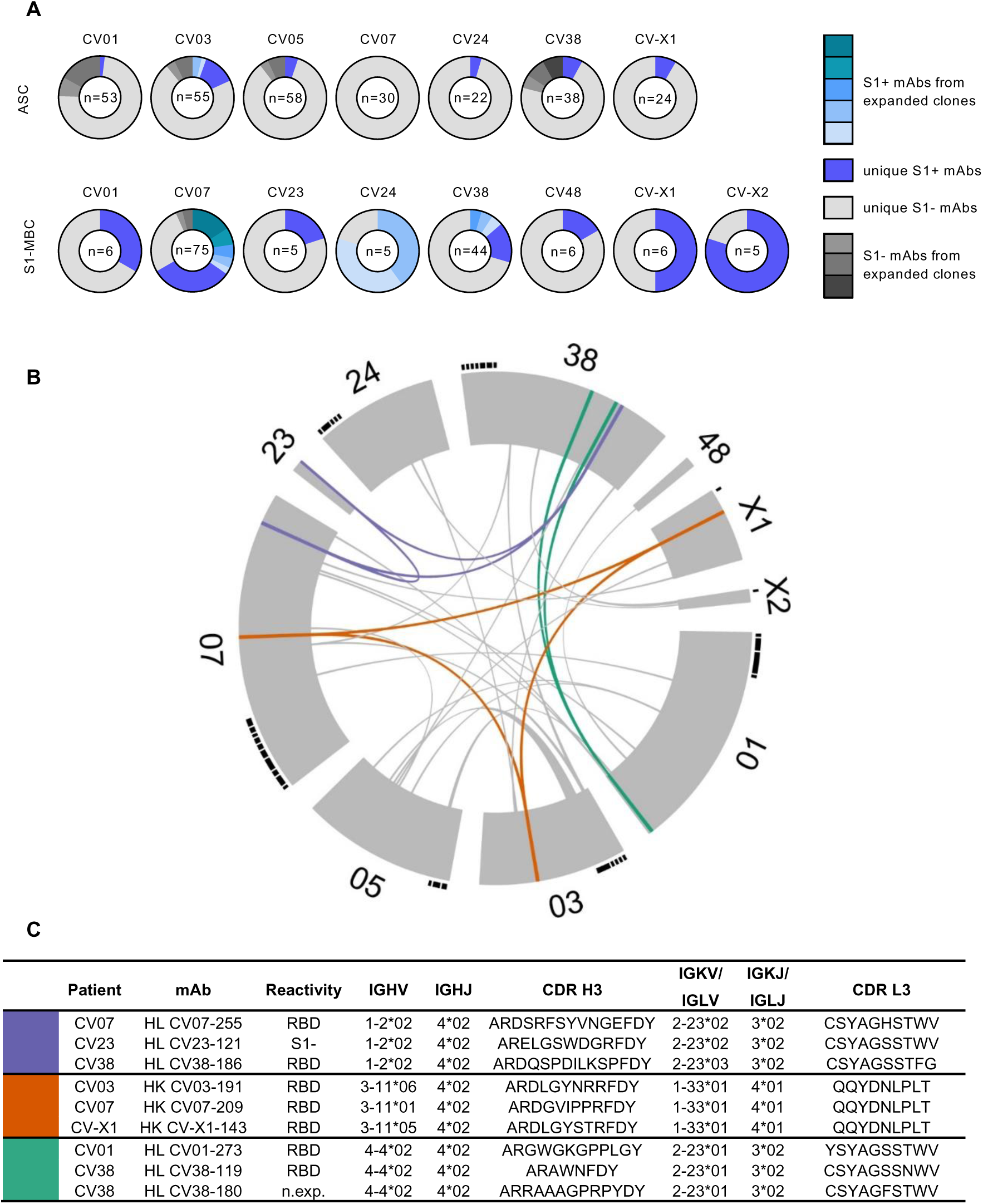
Clonal expansion and public clonotypes. (**A**) Pie charts represent clonal relationship of all expressed mAbs from each donor separately for antibody secreting cells (ASC) and S1- stained memory B cells (S1-MBC). mAbs were considered S1-reactive (S1+) or non-S1-reactive (S1-) based on SARS-CoV-2-S1 ELISA measurements. Antibodies were considered to be clonally expanded when they were isolated from multiple cells. (**B**) Circos plot displays all isolated mAbs from ten donors. Interconnecting lines indicate relationship between mAbs that share the same V and J gene on both Ig heavy and light chain. Such public or shared clonotypes in which more than 50% of mAbs are S1-reactive are represented as colored lines. Small black angles at the outer circle border indicate expanded clones within the respective donor. (**C**) Properties of public clonotypes from S1+ mAbs according to the colors used in (B) with sequence similarities between mAbs isolated from different donors, also within complementarity-determining region (CDR) three. IGHV, IGHJ IGKV, IGKJ, IGLV, IGLJ = V and J genes of immunoglobulin heavy, kappa, lambda chains; n.exp. = not expressed.

**Fig. S5.**
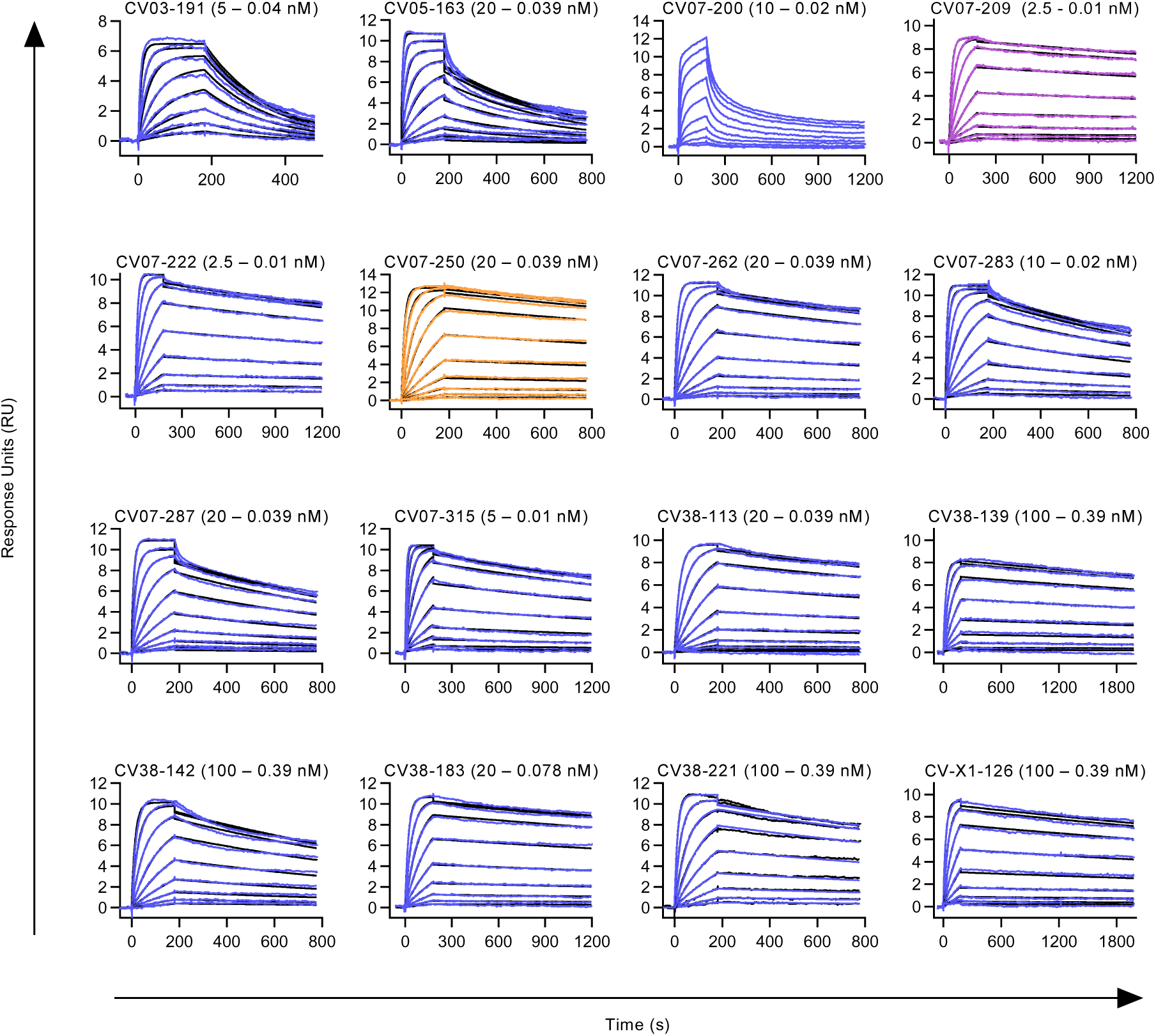
Binding kinetic measurements of mAbs to RBD. Binding kinetics of mAbs to RBD were modeled (black) from multi-cycle surface plasmon resonance (SPR) measurements (blue, purple, orange). Fitted monovalent analyte model is shown. For CV07-200, neither a bivalent nor a monovalent analyte model described the data accurately (no model is shown). Three out of the 18 selected mAbs for detailed characterization (Top-18) were not analyzed using multi-cycle-kinetics: CV07-270 was excluded as it interacted with the anti-mouse IgG reference surface on initial qualitative measurements. CV07-255 and CV-X2-106 were not analyzed since they showed biphasic binding kinetics and relatively fast dissociation rates in initial qualitative measurements. Non-neutralizing CV03-191, a mAb not included in the Top-18 mAbs, was included in the multicycle experiments as it has the same clonotype as strongly neutralizing CV07-209 (Fig. S4C). All measurements are performed by using a serial 2-fold dilution of mAbs on reversibly immobilized SARS-CoV-2-S1 RBD-mFc.

**Fig. S6.**
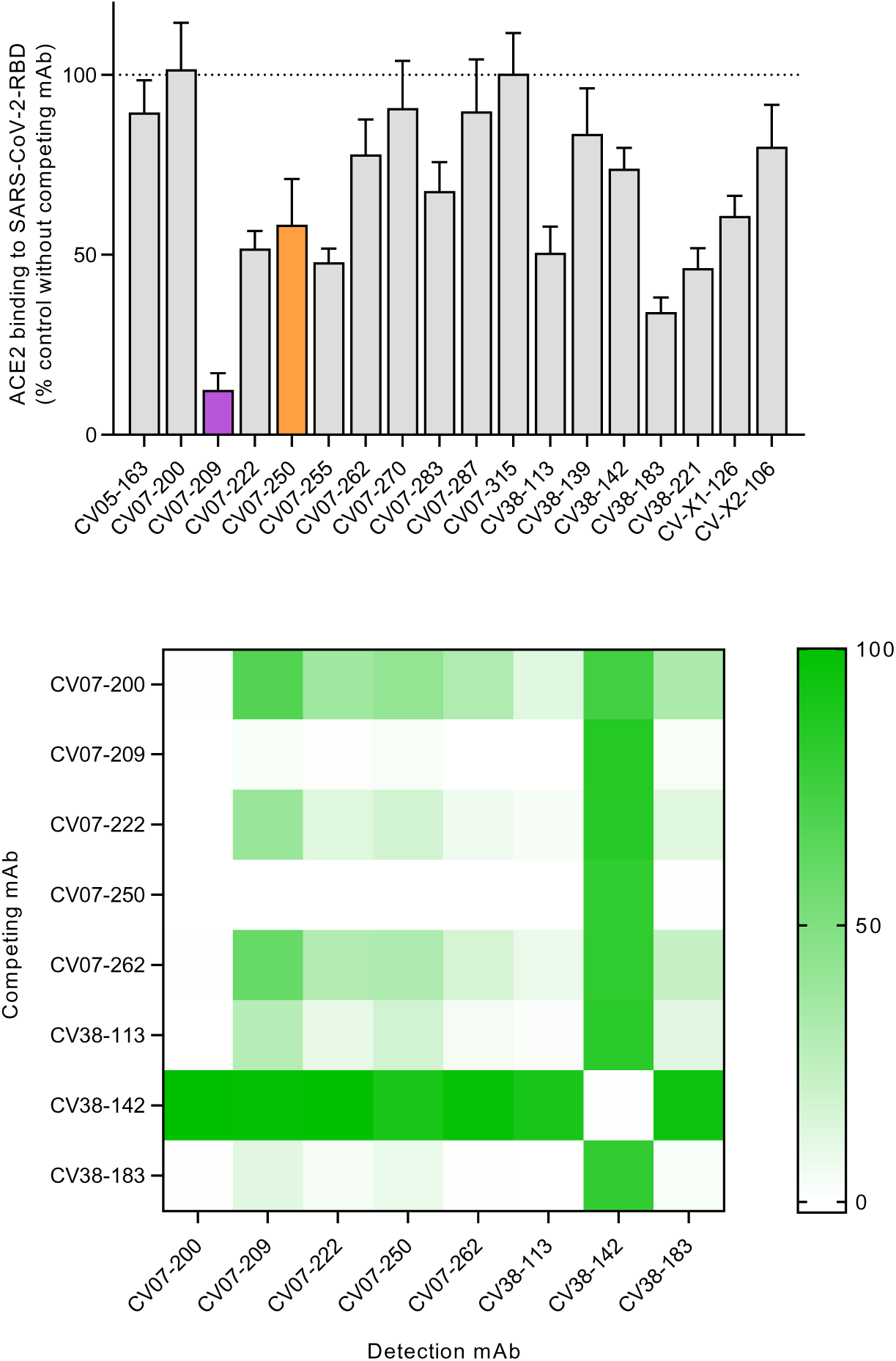
Competition of mAbs for RBD binding with ACE2 and between SARS-CoV-2 mAbs. (**A**) ELISA-based measurements of human ACE2 binding to SARS-CoV-2 RBD after pre-incubation with the indicated neutralizing mAbs. Values are shown relative to antibody-free condition as mean+SD from three independent measurements. (**B**) Competition for RBD binding between combinations of potent neutralizing mAbs is illustrated as a heat map. Shades of green indicate the degree of competition for RBD binding of detection mAb in presence of 100-fold excess of competing mAb relative to non-competition conditions. Green squares indicate no competition. Values are shown as mean of two independent experiments.

**Fig. S7.**
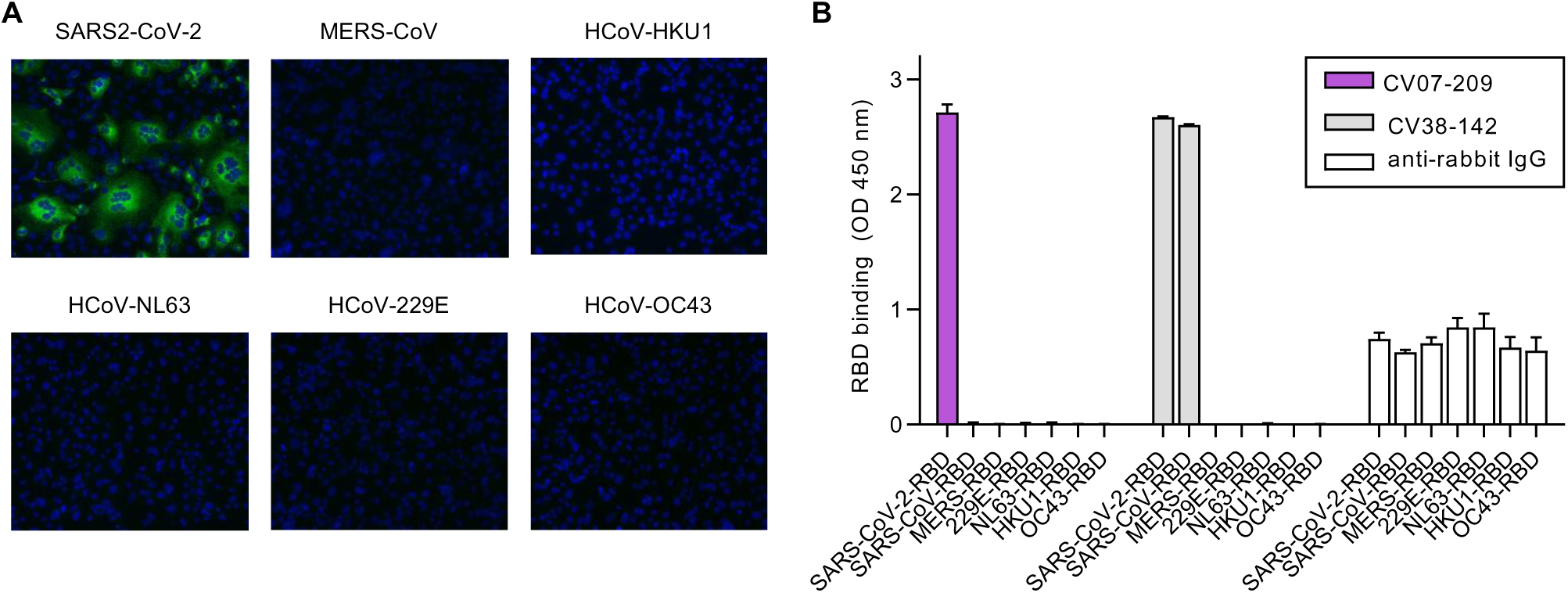
Binding specificity of mAbs within coronaviruses. (**A**) Representative immunofluorescence staining on VeroB4 cells overexpressing spike protein of indicated coronavirus with SARS-CoV-2 mAb CV07-209 at 5 µg/ml. For all other 17 of the selected 18 mAbs (Top-18, Supplementary Table ST4), similar results were obtained. (**B**) Binding of indicated mAbs to fusion proteins containing the RBD of indicated coronaviruses and the constant region of rabbit IgG revealed by ELISA. For all other Top-18 mAbs, similar results were obtained as for CV07-209. Values indicate mean+SD from two wells of one experiment.

**Fig. S8.**
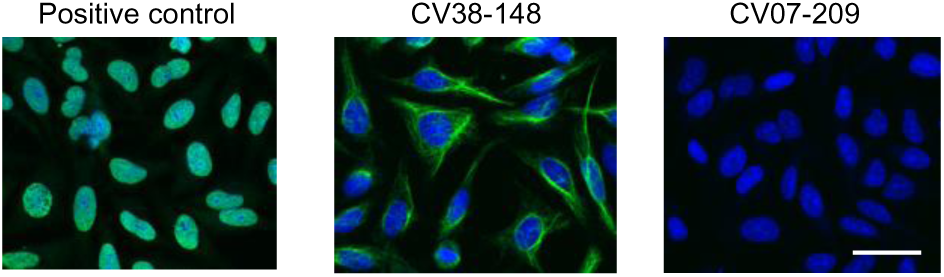
Representative staining on HEp-2 cells of selected S1-reactive antibodies. Representative HEp-2 cell staining with a commercial anti-nuclear antibody as positive control revealed nuclear binding (left). S1-reactive non-neutralizing mAb CV38-148 exhibited cytoplasmatic binding (middle). Neutralizing mAb CV07-209 showed no binding (right). All mAbs selected for detailed characterization (Top-18, Supplementary Table ST4) revealed similar results like CV07-209 when used at 50 µg/ml.

**Fig. S9.**
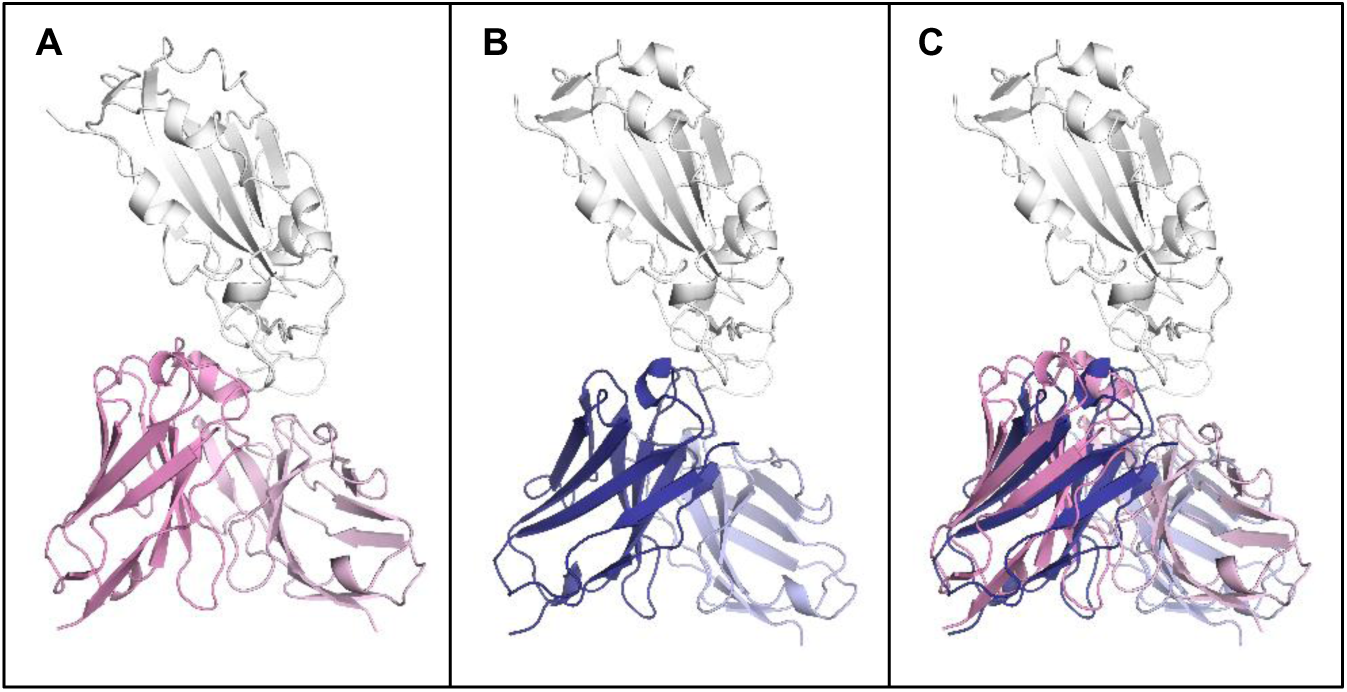
Structural comparison of CV07-270/RBD and P2B-2F6/RBD complexes. (**A**) Structure of CV07-270 (pink) in complex with RBD (white). (**B**) Structure of P2B-2F6 (blue) in complex with RBD (white) (PDB 7BWJ) (Ju et al., 2020) (**C**) Structures of CV07-270/RBD and P2B-2F6/RBD were superimposed based on the RBD.

**Fig. S10.**
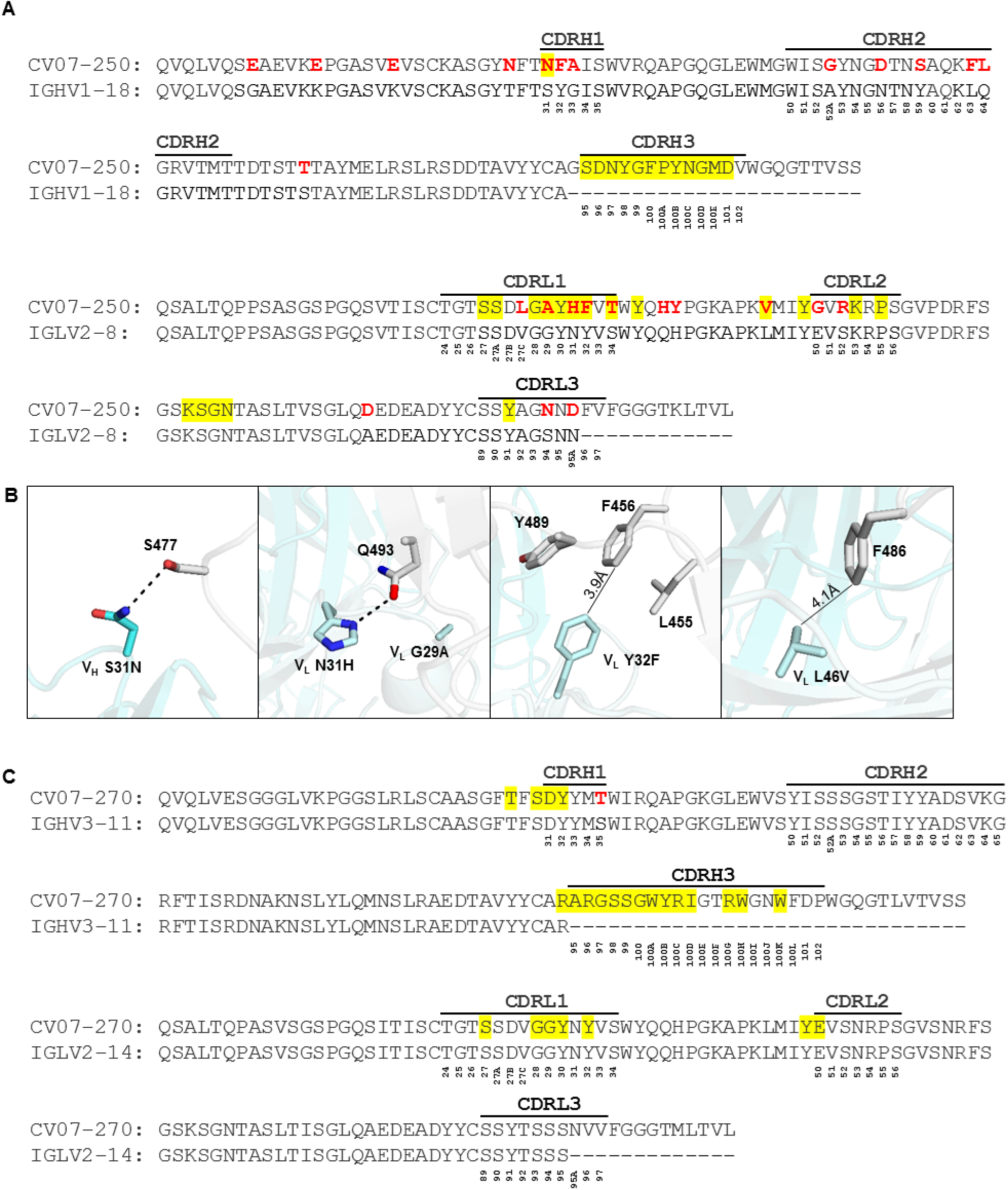
Comparison of sequences of CV07-250 and CV07-270 to their putative germline sequences. (A) Alignment of CV07-250 with the germline IGHV1-18 sequence (nucleotide SHM rate 5.8%) and IGLV2-8 (nucleotide SHM rate 5.4%). (B) Somatic mutations VH S31H, VL G29A, VL N31H, VL Y32F, VL S34T, and VL L46V are located in the CV07-250 paratope with other somatic mutations in all of the CDRs that may affect overall CDR conformation and interactions. Hydrogen bonds are represented by dashed lines. Distances between atoms are shown in solid lines. CV07-250 heavy chain is in dark cyan and light chain is in light cyan. SARS-CoV-2 RBD is in light grey. (**C**) Alignment of CV07-270 with the germline IGHV3-11 sequence (nucleotide SHM rate 0.7%) and IGLV2-14 (nucleotide SHM rate 0%). The regions that correspond to CDR H1, H2, H3, L1, L2, and L3 are indicated. Residues that differ from the germline are highlighted in red. Residues that interact with the RBD are highlighted in yellow. Residue positions in the CDRs are labeled according to the Kabat numbering scheme.

**Supplementary Table ST1.**
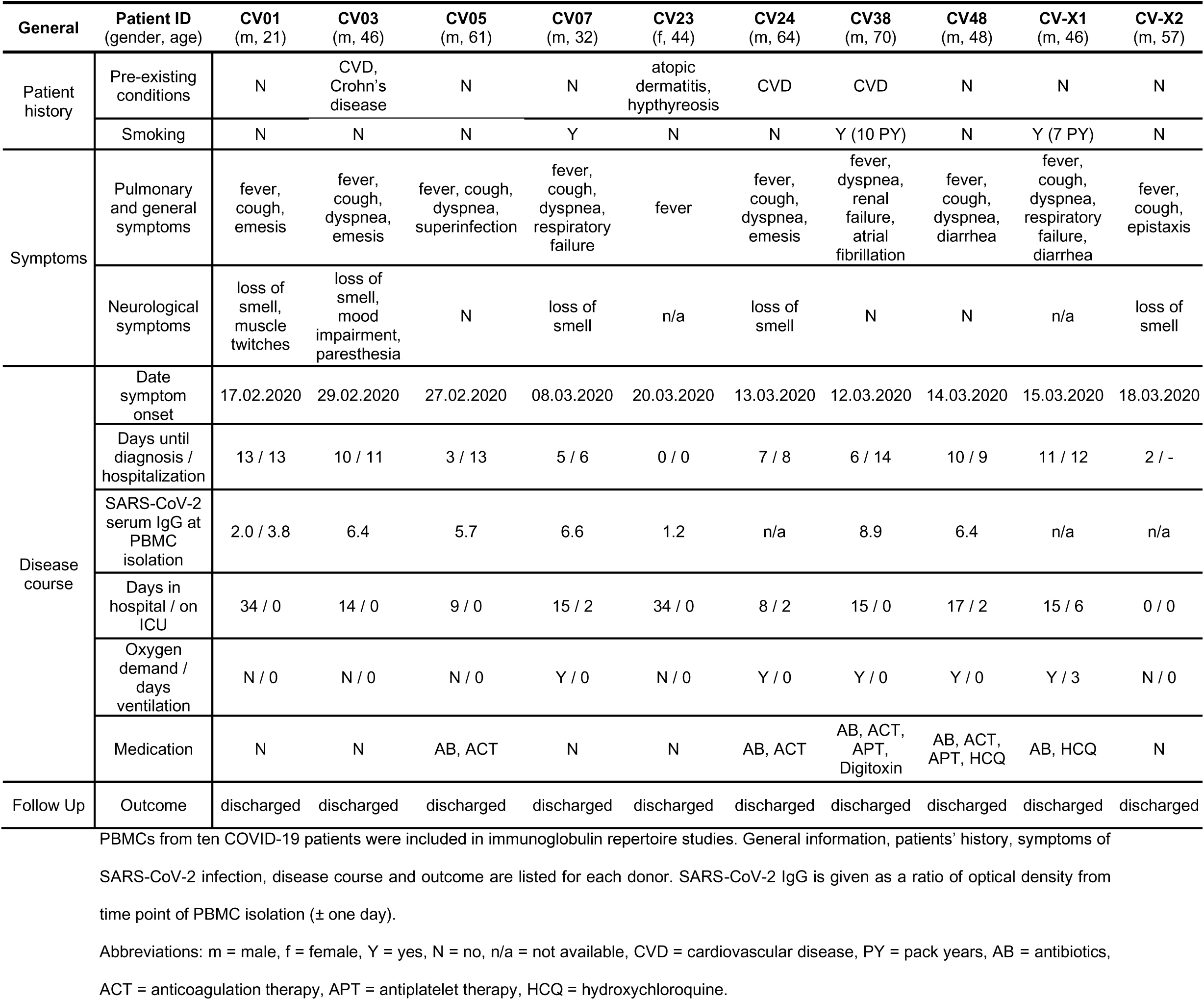
Description of patient cohort.

**Supplementary Table ST2.**
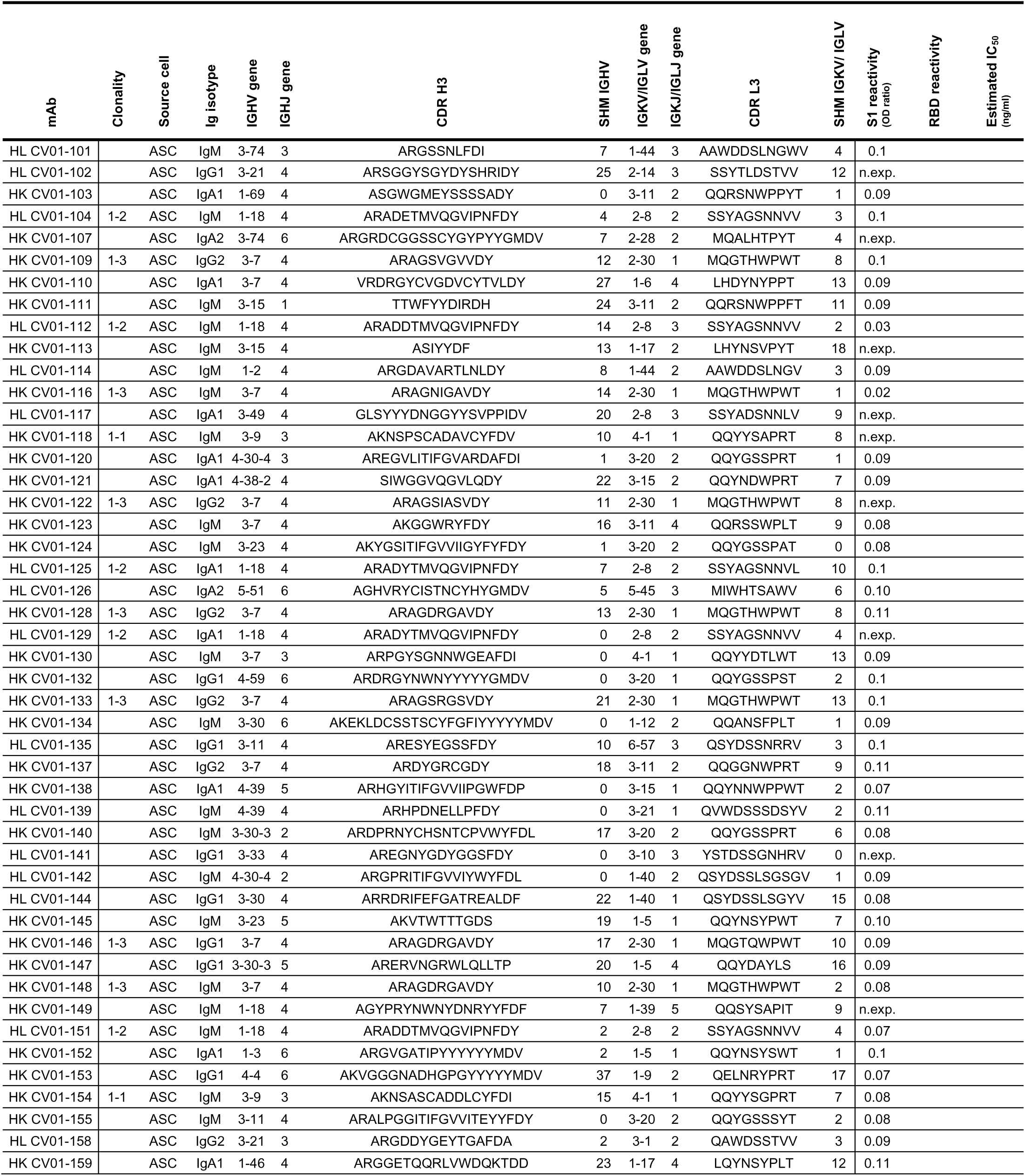
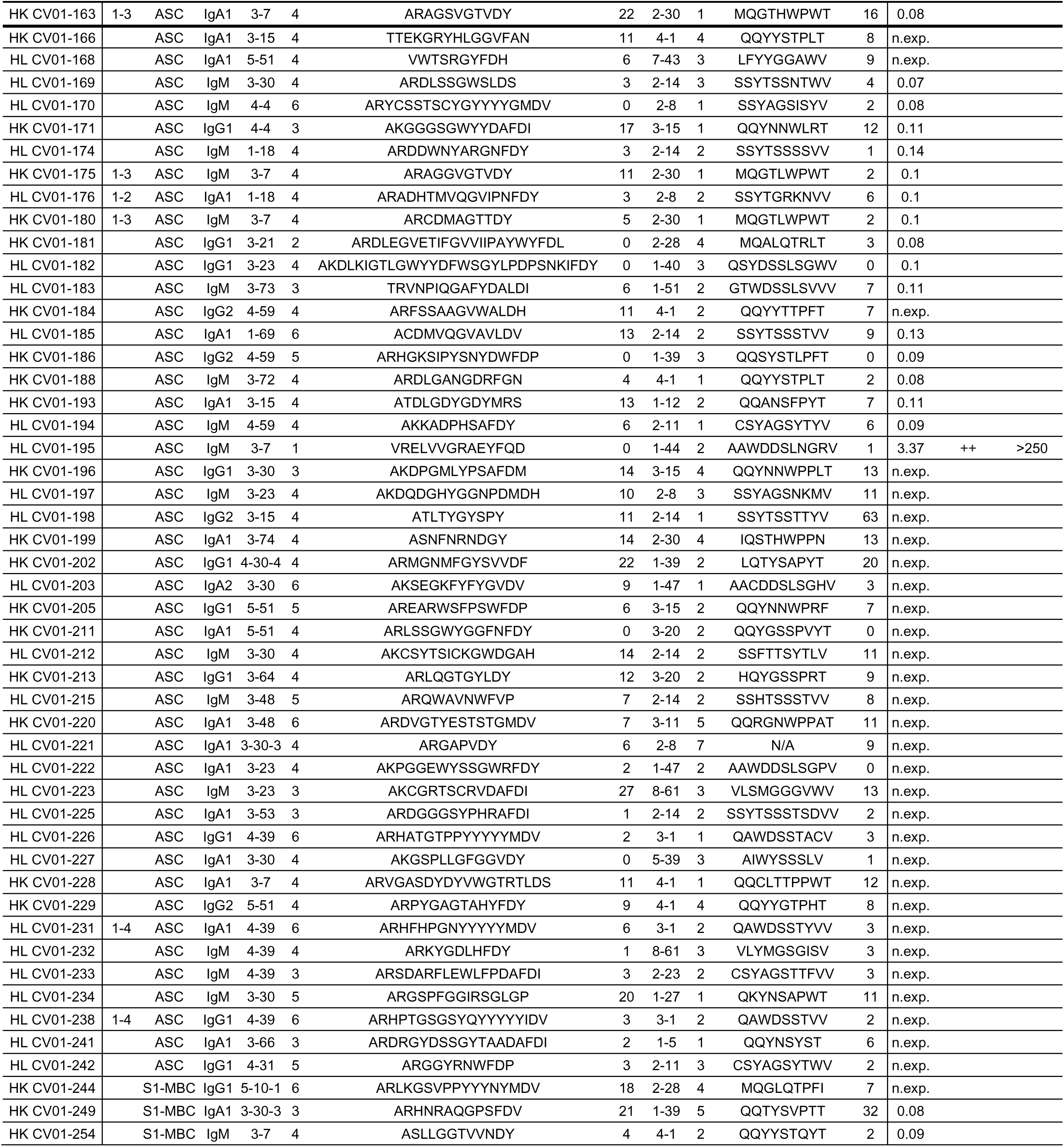
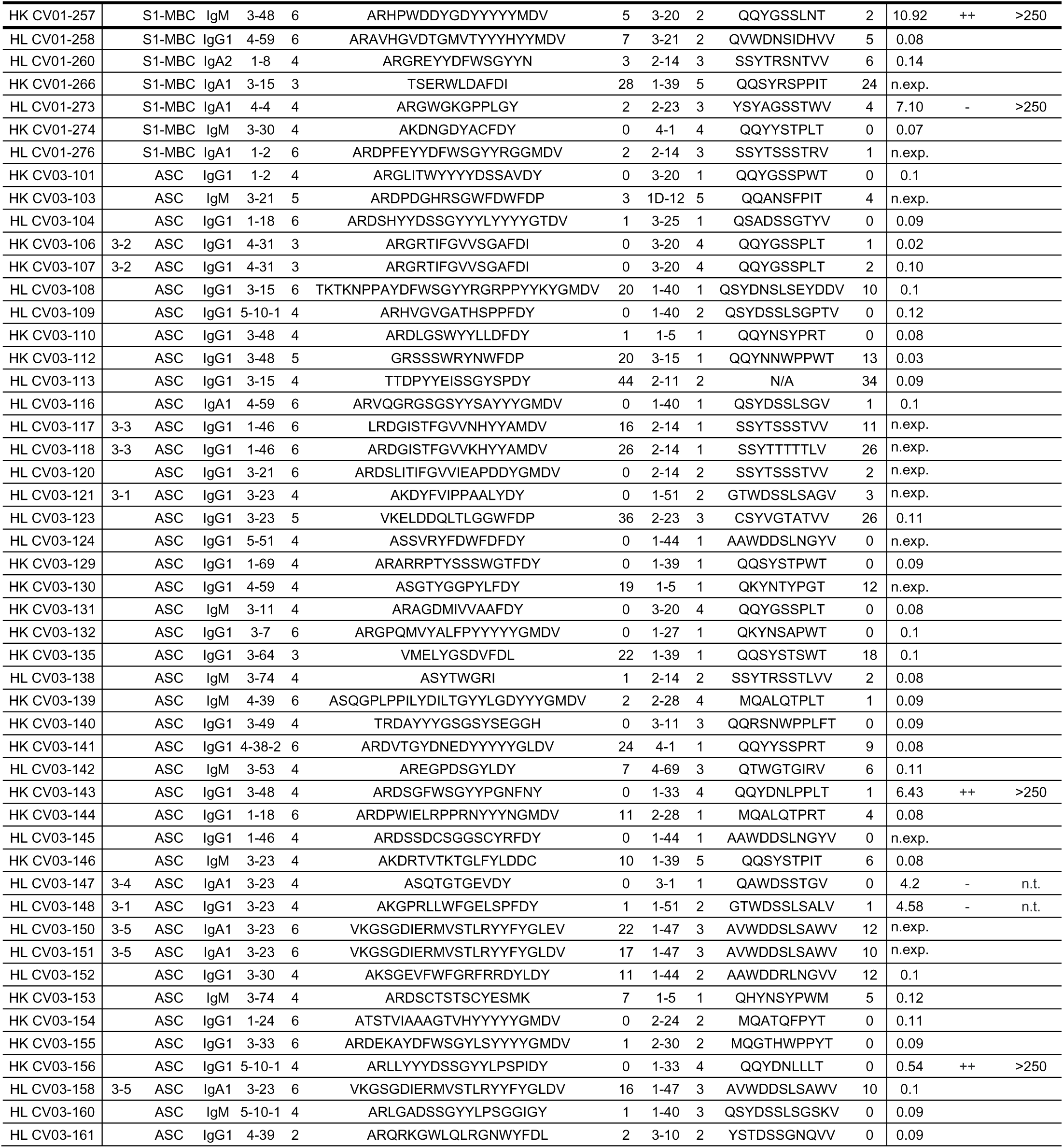
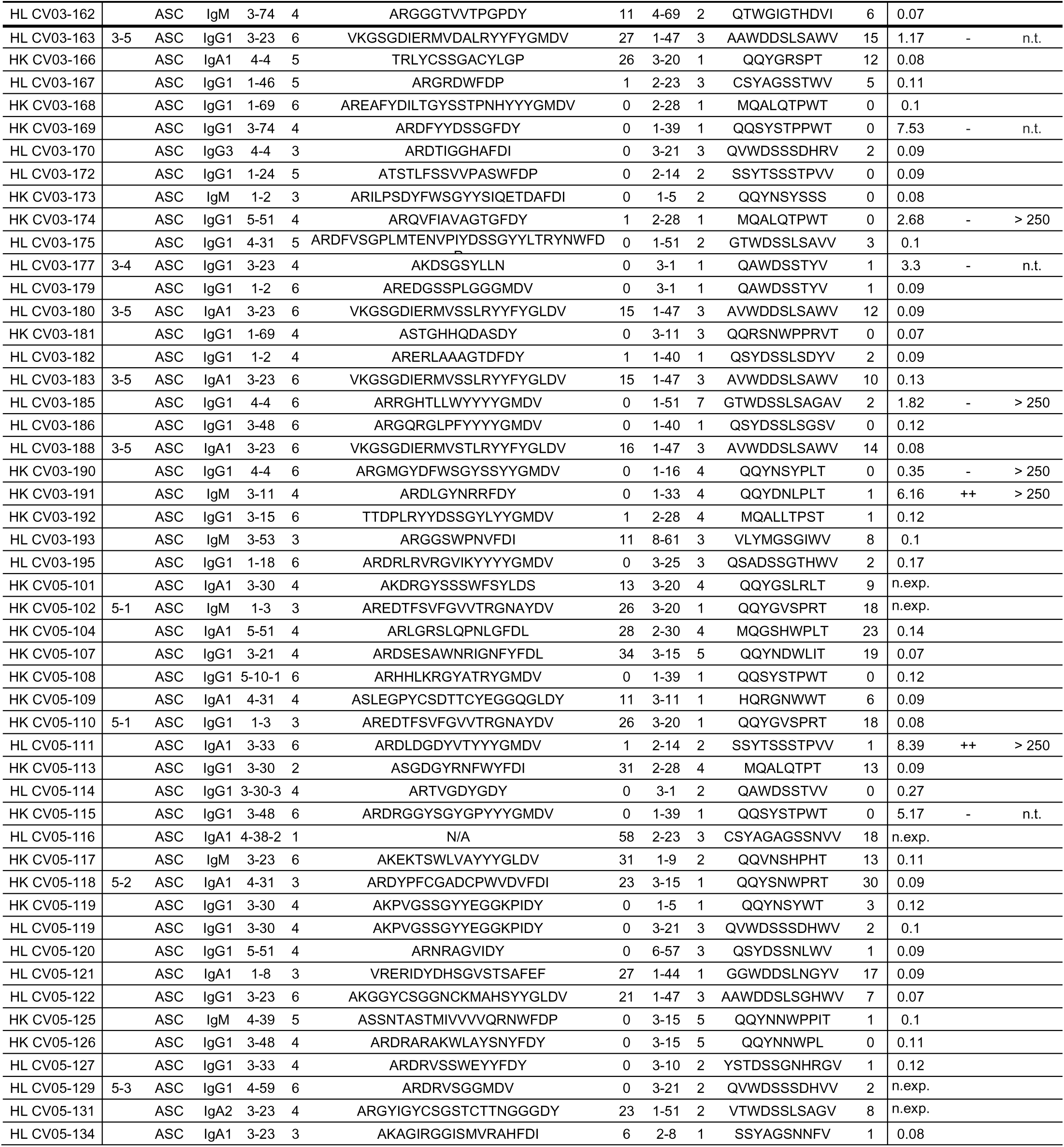
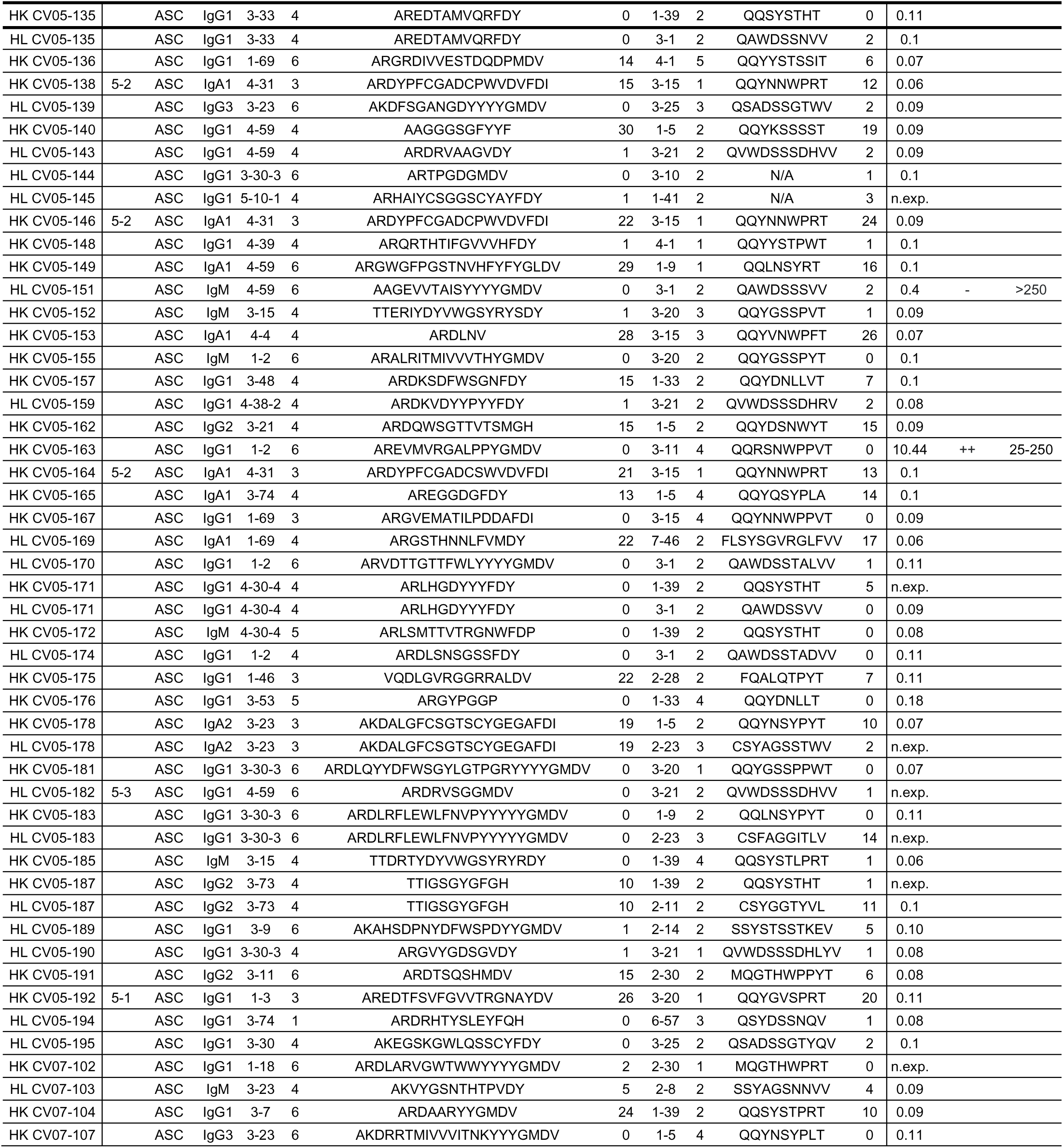
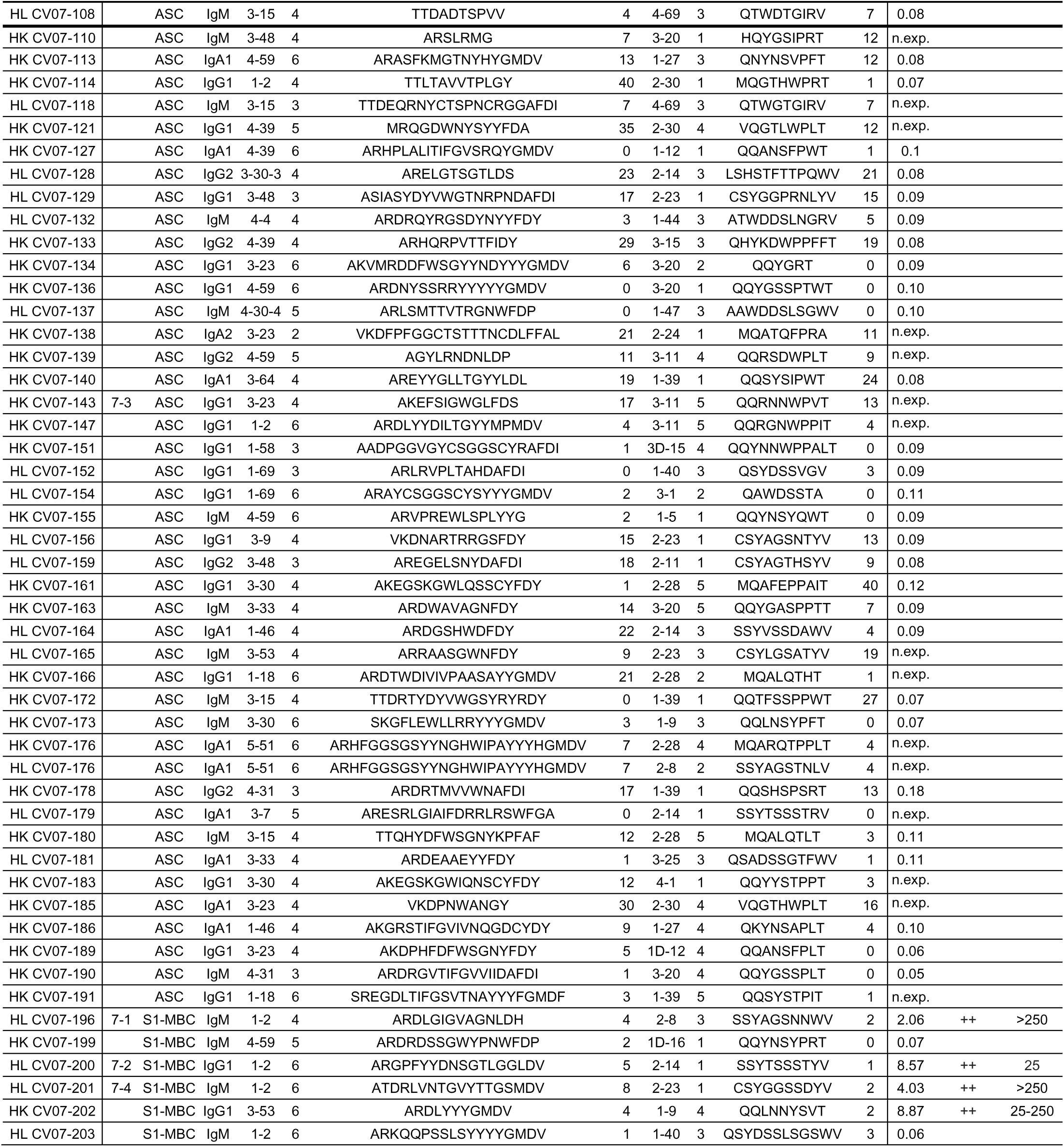
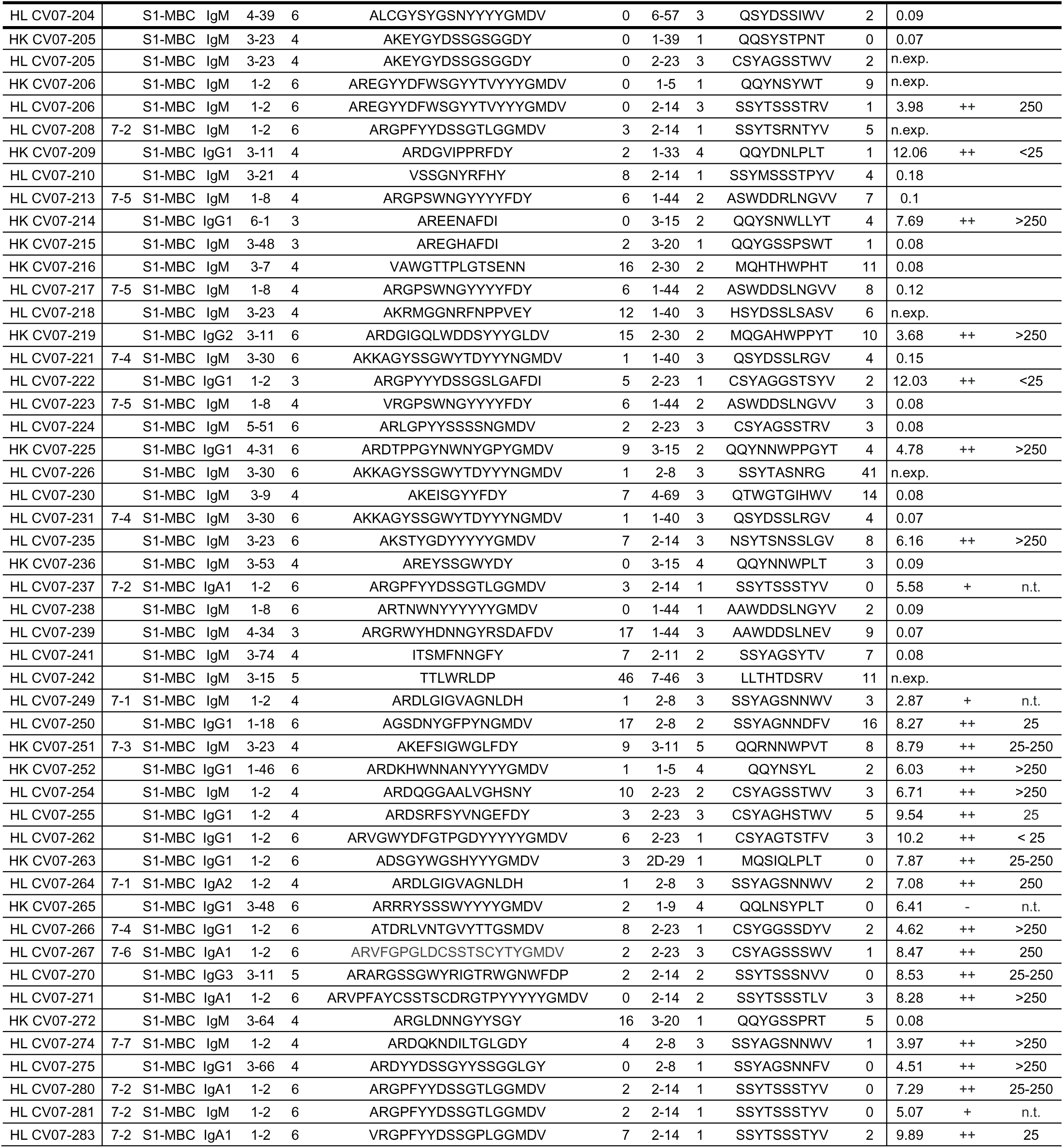
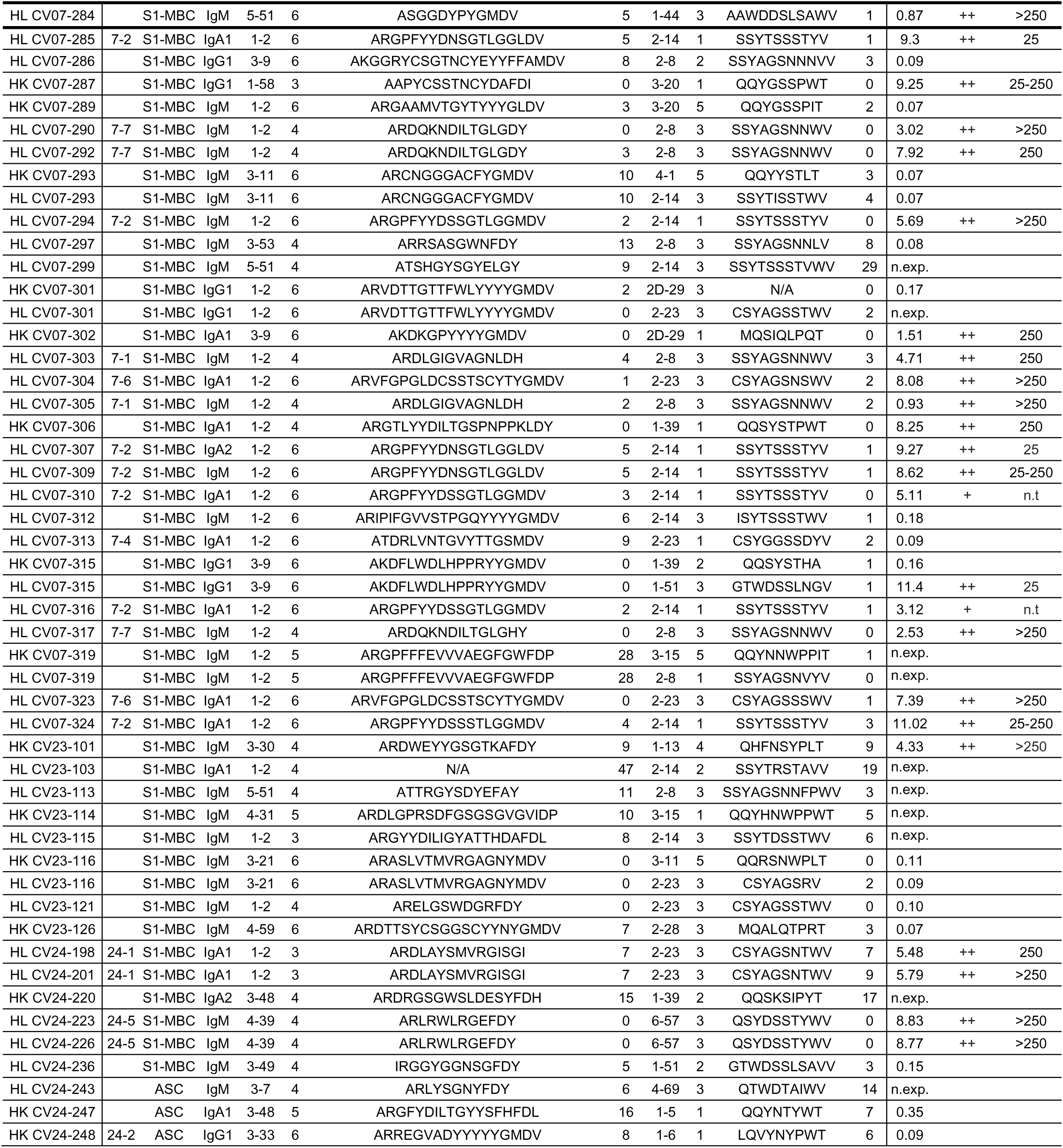
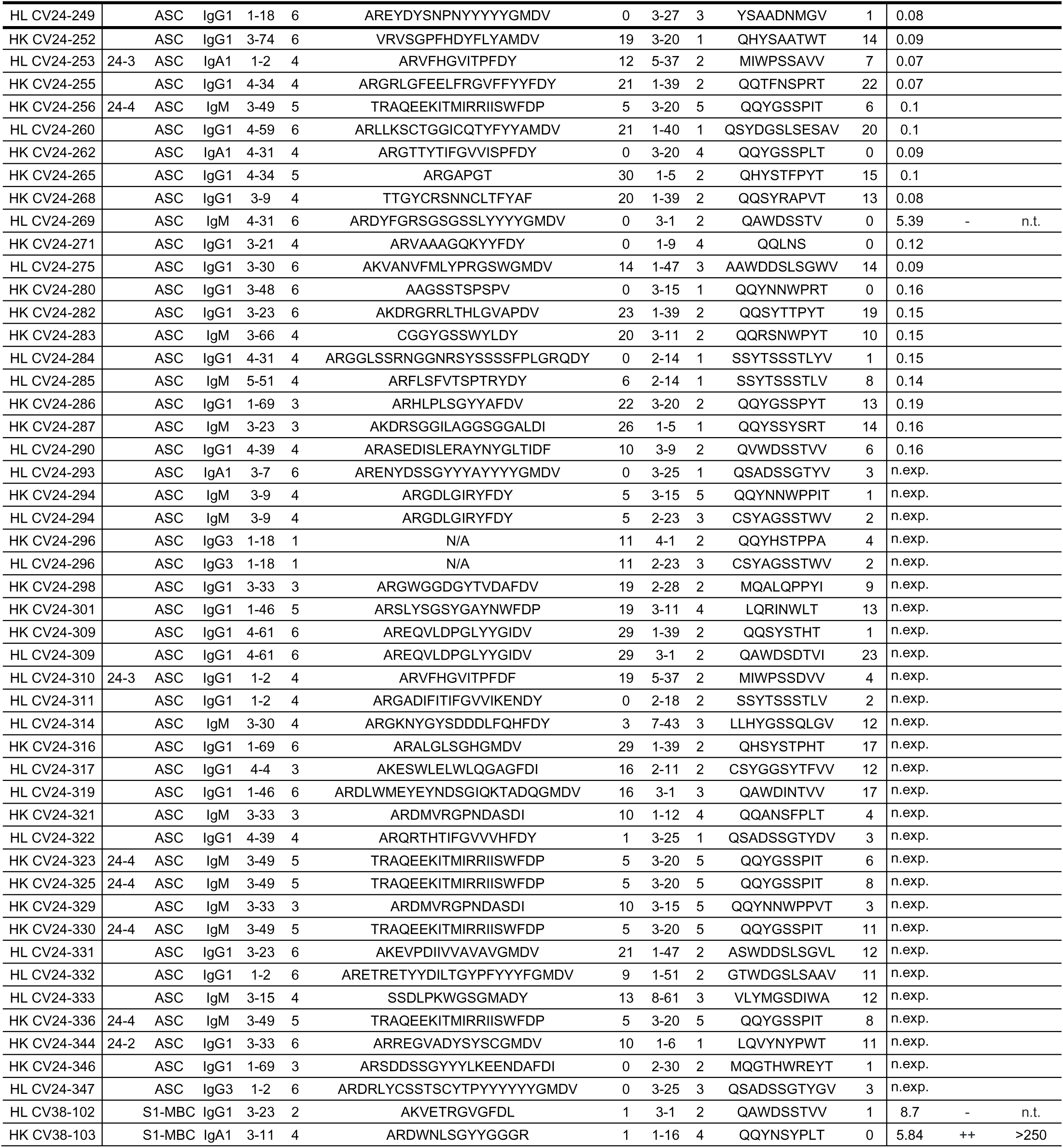
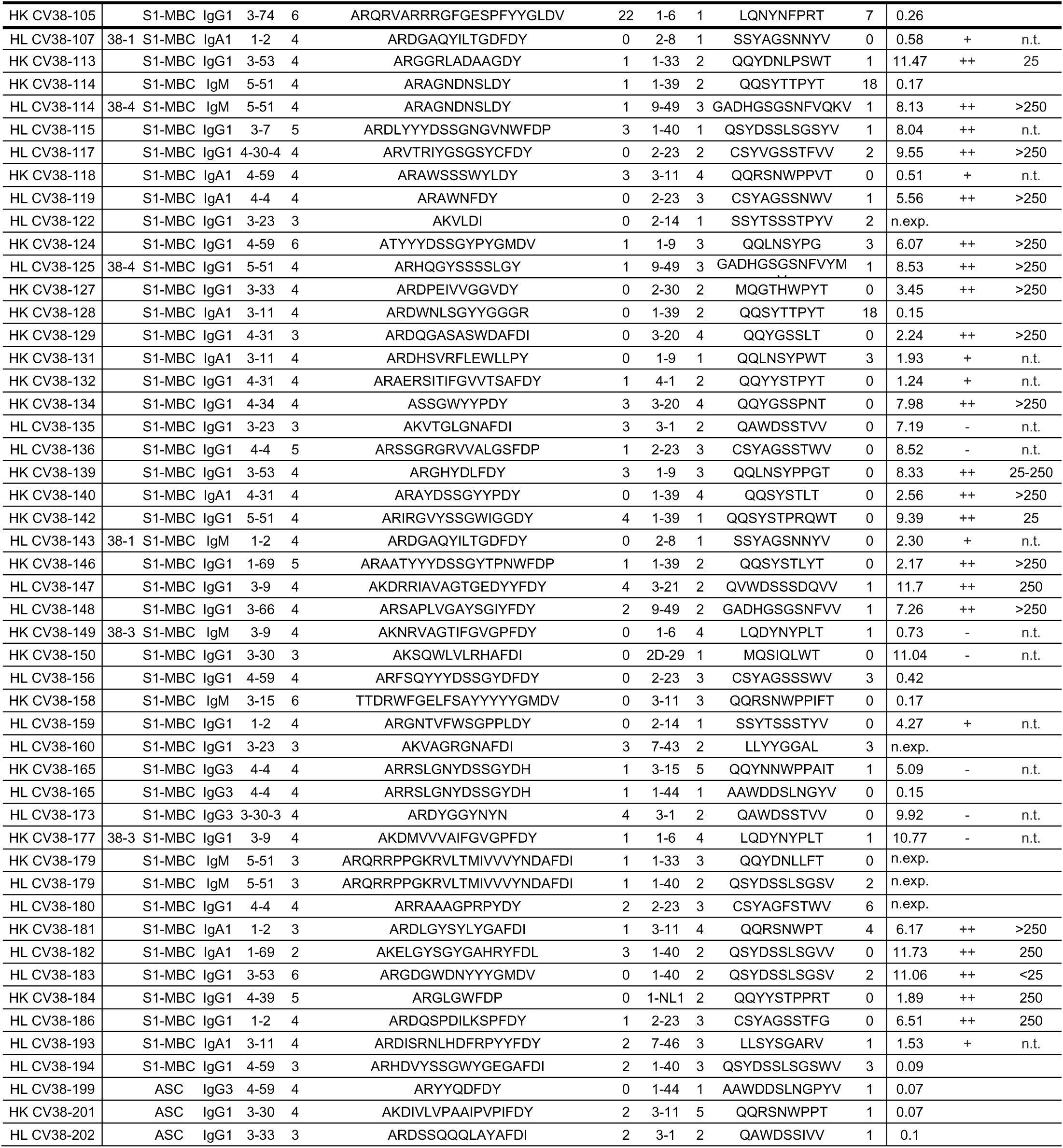
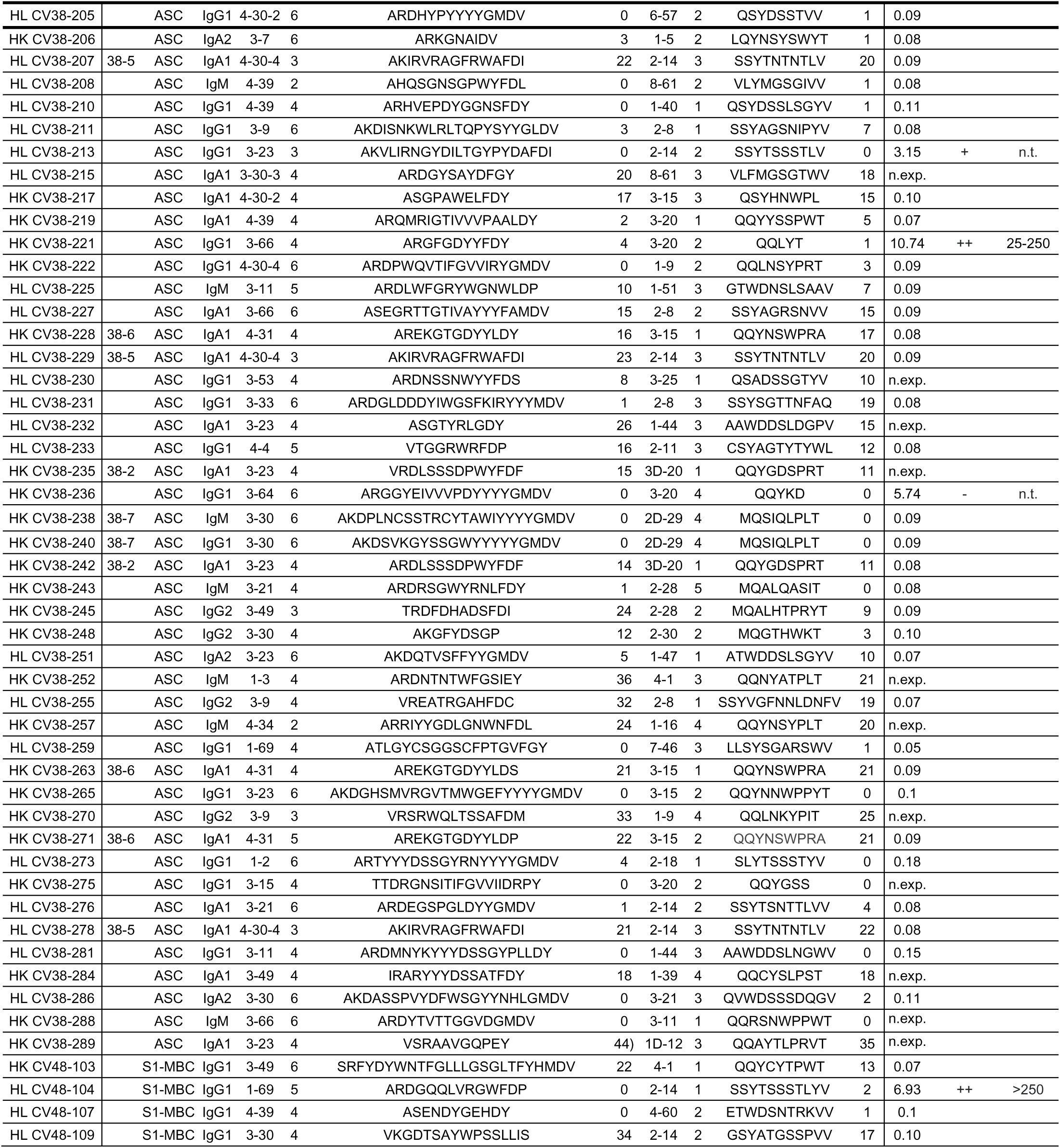
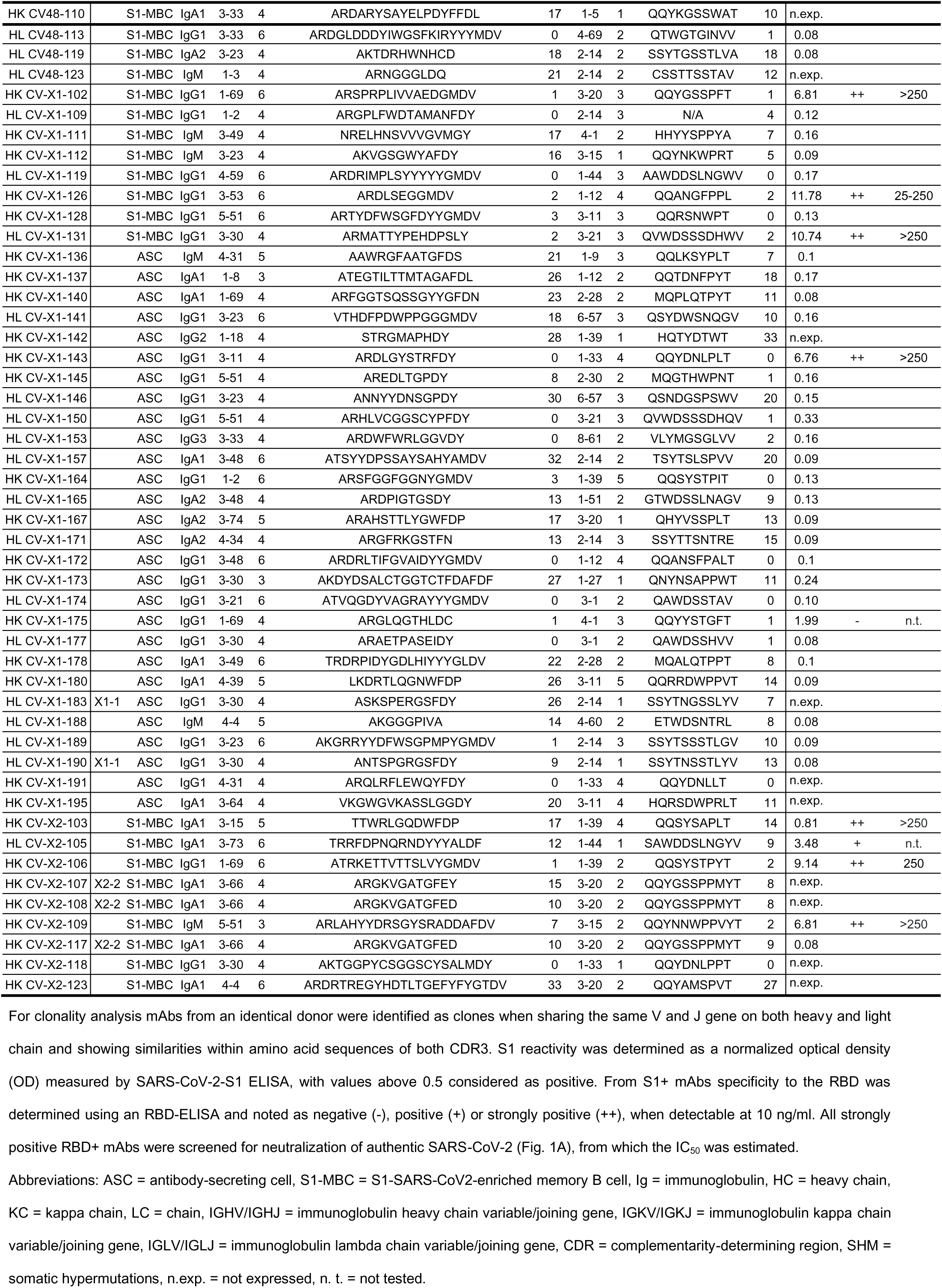
Immunoglobulin sequence and functional screening data of all isolated mAbs.

**Supplementary Table ST3.** Public or common antibody response using VH3-53 and VH3-66 genes.

| mAb | IGHV | IGHJ | CDR H3 | IGKV/IGLV | IGKJ/IGLJ | CDR L3 |
| --- | --- | --- | --- | --- | --- | --- |
| RBD-494 (Cao et al.) | 3-53 | 6 | ARDLVVYGM DV | 1-9 | 4 | QQLNSYPFT |
| HK CV07-202 | 3-53*01 | 6*02 | ARDLYYYGM DV | 1-9*01 | 4*01 | QQLNNYSVT |
| HL CV07-275 | 3-66*01 | 4*02 | ARDYYDSSGYSSGGLGY | 2-8*01 | 1*01 | SSYAGSNNFV |
| HL CV38-148 | 3-66*01 | 4*02 | ARSAPLVGAYSGIYFDY | 9-49*01 | 2*01 | GADHGSGSNFV |
| HK CV38-221 | 3-66*01 | 4*02 | ARGFGDYFDY | 3-20*01 | 2*01 | QQLYT |
| HK CV38-113 | 3-53*04 | 4*02 | ARGGRLADAAGDY | 1-33*01 | 2*02 | QQYDNLPSWT |
| HK CV38-139 | 3-53*01 | 4*02 | ARGHYDLFDY | 1-9*01 | 3*01 | QQLNSYPPGT |
| HL CV38-183 | 3-53*01 | 6*02 | ARGDGWDNYYYGM DV | 1-40*01 | 2*01 | QSYDSSLGSGV |
| HK CV-X1-126 | 3-53*04 | 6*02 | ARDLSEGGMDV | 1-12*01 | 4*01 | QQANGFSAL |

Abbreviations: IGHV/IGHJ = immunoglobulin heavy chain variable/joining gene, IGKV/IGKJ = immunoglobulin kappa chain variable/joining gene. IGLV/IGLJ = immunoglobulin lambda chain variable/joining gene, CDR = complementarity-determining region.

**Supplementary Table ST4.** Biophysical and functional characterization of the 18 most potent SARS-CoV-2 neutralizing mAbs.

| mAb | EC<sub>50</sub><br>(ng/ml) | K<sub>a</sub><br>(s<sup>-1</sup>) | K<sub>d</sub><br>(M<sup>-1</sup>s<sup>-1</sup>) | K<sub>D</sub><br>(M) | IC<sub>50</sub><br>(ng/ml) | ACE2 binding<br>reduction (%) | Tissue<br>reactivity |
| --- | --- | --- | --- | --- | --- | --- | --- |
| CV05-163 | 3.8 | 8.48 E+06 | 1.86 E-03 | 2.20 E-10 | 16.3 | 9.6 | none |
| CV07-200 | 9.0 | n/a | n/a | n/a | 14.5 | none | B / L* |
| CV07-209 | 4.1 | 2.40 E+07 | 1.44 E-04 | 6.00 E-12 | 3.1 | 73.9 | none |
| CV07-222 | 5.5 | 2.98 E+07 | 2.07 E-04 | 6.97 E-12 | 7.8 | 36.9 | B / L / C |
| CV07-250 | 8.7 | 4.05 E+06 | 2.28 E-04 | 5.64 E-11 | 3.5 | 59.9 | none |
| CV07-255 | 4.8 | n/a | n/a | n/a | 14.5 | 51.6 | L / H / C / K |
| CV07-262 | 12.1 | 4.27 E+06 | 3.37 E-04 | 7.90 E-11 | 7.1 | 25.5 | none |
| CV07-270 | 14.2 | n/a | n/a | n/a | 82.3 | 7.5 | L / H* / C |
| CV07-283 | 4.0 | 1.52 E+07 | 7.24 E-04 | 4.75 E-11 | 16.9 | 38.0 | none |
| CV07-287 | 5.9 | 5.11 E+06 | 8.14 E-04 | 1.60 E-10 | 41.7 | 12.4 | none |
| CV07-315 | 7.2 | 2.36 E+07 | 2.62 E-04 | 1.12 E-11 | 24.9 | none | none |
| CV38-113 | 5.2 | 2.33 E+06 | 3.12 E-04 | 1.34 E-10 | 20.8 | 51.2 | none |
| CV38-139 | 8.9 | 3.62 E+05 | 1.00 E-04 | 2.77 E-10 | 73.2 | 14.9 | none |
| CV38-142 | 5.6 | 6.31 E+05 | 6.60 E-04 | 1.05 E-09 | 23.2 | 26.9 | none |
| CV38-183 | 5.4 | 2.31 E+06 | 1.37 E-04 | 5.93 E-11 | 3.7 | 65.9 | none |
| CV38-221 | 6.4 | 7.45 E+05 | 3.71 E-04 | 4.99 E-10 | 172.6 | 50.1 | none |
| CV-X1-126 | 5.5 | 4.12 E+05 | 9.98 E-05 | 2.43 E-10 | 71.7 | 44.2 | none |
| CV-X2-106 | 5.6 | n/a | n/a | n/a | 17.6 | 19.9 | none |

Abbreviations: EC<sub>50</sub> = half-maximal effective concentration, K<sub>a</sub> = association rate, K<sub>d</sub> = dissociation rate, K<sub>D</sub> = equilibrium dissociation constant, IC<sub>50</sub> = half-maximal inhibitory concentration, B = brain, L = lung, H = heart, C = colon, K = kidney. An asterisk (\*) indicates weak binding.

**Supplementary Table ST5.**
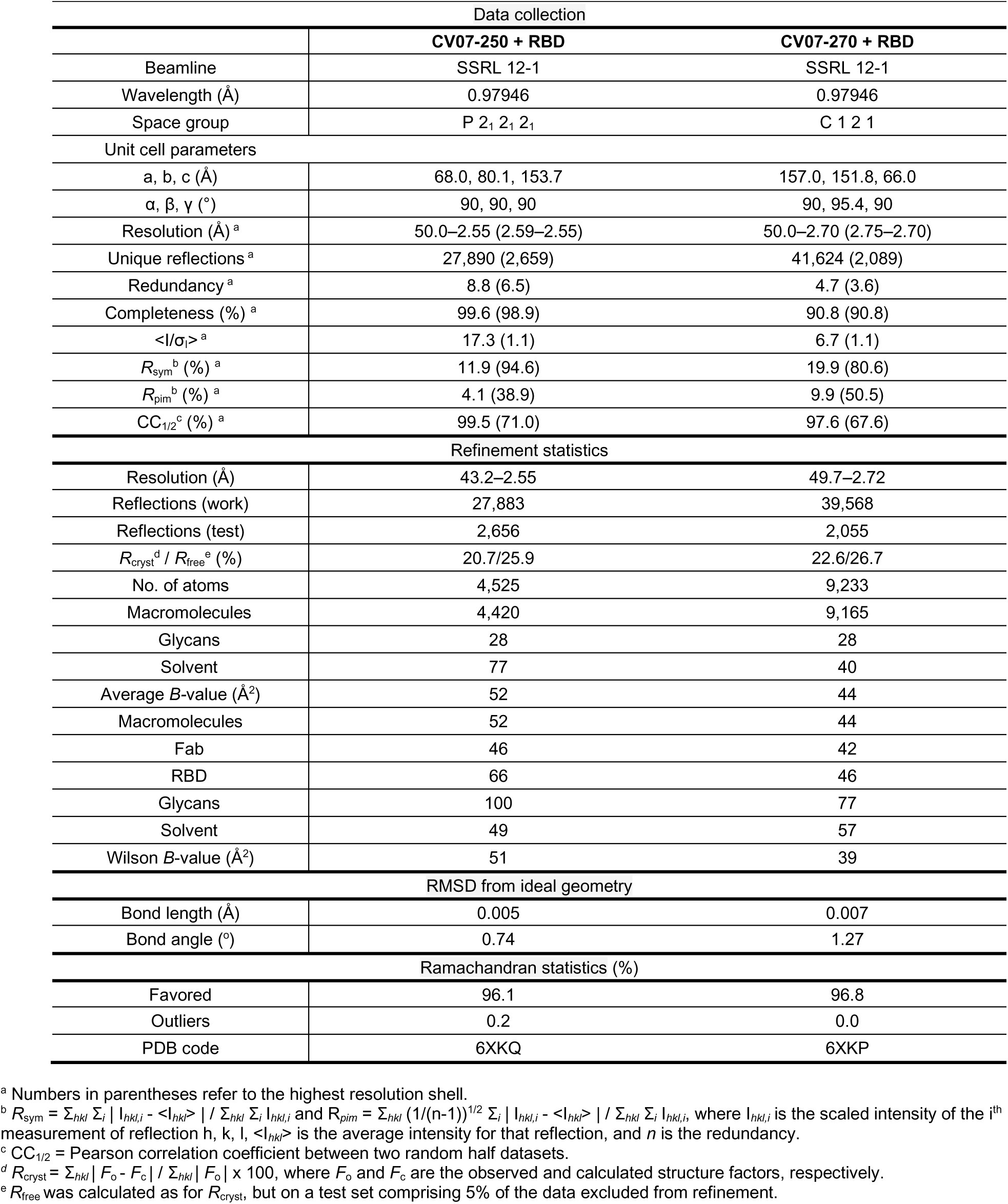
X-ray data collection and refinement statistics.

| Data collection |  |  |
| --- | --- | --- |
|  | CV07-250 + RBD | CV07-270 + RBD |
| Beamline | SSRL 12-1 | SSRL 12-1 |
| Wavelength (Å) | 0.97946 | 0.97946 |
| Space group | P 2<sub>1</sub> 2<sub>1</sub> 2<sub>1</sub> | C 1 2 1 |
| Unit cell parameters |  |  |
| a, b, c (Å) | 68.0, 80.1, 153.7 | 157.0, 151.8, 66.0 |
| α, β, γ (°) | 90, 90, 90 | 90, 95.4, 90 |
| Resolution (Å)<sup>a</sup> | 50.0–2.55 (2.59–2.55) | 50.0–2.70 (2.75–2.70) |
| Unique reflections<sup>a</sup> | 27,890 (2,659) | 41,624 (2,089) |
| Redundancy<sup>a</sup> | 8.8 (6.5) | 4.7 (3.6) |
| Completeness (%)<sup>a</sup> | 99.6 (98.9) | 90.8 (90.8) |
| <I/σ<sub>i</sub>><sup>a</sup> | 17.3 (1.1) | 6.7 (1.1) |
| R<sub>sym</sub><sup>b</sup> (%)<sup>a</sup> | 11.9 (94.6) | 19.9 (80.6) |
| R<sub>pim</sub><sup>b</sup> (%)<sup>a</sup> | 4.1 (38.9) | 9.9 (50.5) |
| CC<sub>1/2</sub><sup>c</sup> (%)<sup>a</sup> | 99.5 (71.0) | 97.6 (67.6) |
| Refinement statistics |  |  |
| Resolution (Å) | 43.2–2.55 | 49.7–2.72 |
| Reflections (work) | 27,883 | 39,568 |
| Reflections (test) | 2,656 | 2,055 |
| R<sub>cryst</sub><sup>d</sup> / R<sub>free</sub><sup>e</sup> (%) | 20.7/25.9 | 22.6/26.7 |
| No. of atoms | 4,525 | 9,233 |
| Macromolecules | 4,420 | 9,165 |
| Glycans | 28 | 28 |
| Solvent | 77 | 40 |
| Average B-value (Å<sup>2</sup>) | 52 | 44 |
| Macromolecules | 52 | 44 |
| Fab | 46 | 42 |
| RBD | 66 | 46 |
| Glycans | 100 | 77 |
| Solvent | 49 | 57 |
| Wilson B-value (Å<sup>2</sup>) | 51 | 39 |
| RMSD from ideal geometry |  |  |
| Bond length (Å) | 0.005 | 0.007 |
| Bond angle (°) | 0.74 | 1.27 |
| Ramachandran statistics (%) |  |  |
| Favored | 96.1 | 96.8 |
| Outliers | 0.2 | 0.0 |
| PDB code | 6XKQ | 6XKP |

<sup>a</sup> Numbers in parentheses refer to the highest resolution shell.
<sup>b</sup> $R_{\text{sym}} = \sum_{hkl} \sum_i |I_{hkl,i} - \langle I_{hkl} \rangle| / \sum_{hkl} \sum_i I_{hkl,i}$ and $R_{\text{pim}} = \sum_{hkl} (1/(n-1))^{1/2} \sum_i |I_{hkl,i} - \langle I_{hkl} \rangle| / \sum_{hkl} \sum_i I_{hkl,i}$ where $I_{hkl,i}$ is the scaled intensity of the $i^{\text{th}}$ measurement of reflection $h, k, l$ , $\langle I_{hkl} \rangle$ is the average intensity for that reflection, and $n$ is the redundancy.
<sup>c</sup> CC<sub>1/2</sub> = Pearson correlation coefficient between two random half datasets.
<sup>d</sup> $R_{\text{cryst}} = \sum_{hkl} |F_o - F_c| / \sum_{hkl} |F_o| \times 100$ , where $F_o$ and $F_c$ are the observed and calculated structure factors, respectively.
<sup>e</sup> $R_{\text{free}}$ was calculated as for $R_{\text{cryst}}$ , but on a test set comprising 5% of the data excluded from refinement.

**Supplementary Table ST6.** Hydrogen bonds and salt bridges identified at the antibody-RBD interface using the PISA program.

| CV07-250 | Distance (Å) | SARS-CoV-2 RBD |
| --- | --- | --- |
| **Hydrogen bonds** |  |  |
| H:TYR98[N] | 3.0 | A:ALA475[O] |
| H:ASN31[ND2] | 3.4 | A:SER477[OG] |
| H:ASN100C[ND2] | 3.8 | A:GLY485[O] |
| H:ASN100C[ND2] | 2.9 | A:PHE486[O] |
| H:ASN97[ND2] | 2.8 | A:ASN487[OD1] |
| H:ASN97[ND2] | 2.7 | A:TYR489[OH] |
| H:TYR100B[N] | 3.5 | A:TYR489[OH] |
| H:ASP96[OD1] | 3.0 | A:SER477[N] |
| H:ASP96[OD2] | 2.7 | A:SER477[OG] |
| H:ASP96[OD2] | 3.5 | A:THR478[N] |
| H:ASP96[OD2] | 2.9 | A:THR478[OG1] |
| H:SER95[OG] | 2.9 | A:ASN487[ND2] |
| H:PHE100[O] | 3.4 | A:TYR489[OH] |
| L:SER27A[OG] | 3.2 | A:TYR505[OH] |
| L:HIS31[N] | 3.1 | A:GLN493[OE1] |
| L:SER67[OG] | 3.5 | A:GLY446[O] |
| L:GLY68[N] | 3.1 | A:GLY446[O] |
| L:GLY68[N] | 3.1 | A:TYR449[OH] |
| L:ASN69[N] | 3.8 | A:GLN498[OE1] |
| L:ASN69[ND2] | 3.7 | A:GLN498[OE1] |
| L:TYR49[OH] | 3.5 | A:PHE486[N] |

| CV07-270 | Distance (Å) | SARS-CoV-2 RBD |
| --- | --- | --- |
| **Hydrogen bonds** |  |  |
| H:TYR100B[OH] | 2.8 | A:SER349[OG] |
| H:SER98[N] | 3.2 | A:GLY447[O] |
| H:SER98[OG] | 3.8 | A:GLY447[O] |
| H:SER98[OG] | 3.2 | A:ASN448[OD1] |
| H:ARG100G[NH2] | 2.9 | A:GLU484[OE1] |
| H:ARG100G[NH1] | 3.2 | A:GLU484[OE2] |
| H:ARG100G[NH2] | 3.0 | A:GLU484[OE2] |
| H:THR28[OG1] | 3.2 | A:LYS444[NZ] |
| H:ASP31[OD1] | 2.7 | A:ARG346[NH1] |
| H:ASP31[OD2] | 2.7 | A:LYS444[NZ] |
| H:SER98[O] | 2.7 | A:TYR449[N] |
| H:SER98[O] | 3.3 | A:ASN450[N] |
| H:TYR100B[OH] | 3.6 | A:SER349[N] |
| **Salt bridges** |  |  |
| H:ARG100G[NH2] | 2.9 | A:GLU484[OE1] |
| H:ARG100G[NH1] | 3.2 | A:GLU484[OE2] |
| H:ARG100G[NH2] | 3.0 | A:GLU484[OE2] |
| H:ASP31[OD1] | 2.7 | A:ARG346[NH1] |
| H:ASP31[OD1] | 3.4 | A:ARG346[NE] |
| H:ASP31[OD2] | 2.7 | A:LYS444[NZ] |

**Supplementary Table ST7.** Histopathological scoring of lung tissue from COVID-19 hamster model.

| Animal number | Time point (dpi) | pneumonia |  |  |  |  | Epithelial cell necrosis of bronchi<sup>a</sup> | Bronchial inflammation<sup>a</sup> | Hyperplasia of bronchial epithelial cells<sup>b</sup> | Alveolar epithelial cell necrosis | Hyperplasia type II alveolar epithelial cells<sup>b</sup> | Alveolar edema<sup>c</sup> | Perivascular edema<sup>c</sup> | Perivascular lymphocytic cuffing | Endotheliitis | <sup>a</sup> Bronchitis score | <sup>b</sup> Regeneration score | <sup>c</sup> Edema score |
| --- | --- | --- | --- | --- | --- | --- | --- | --- | --- | --- | --- | --- | --- | --- | --- | --- | --- | --- |
|  |  | % Affected lung tissue | Degree of inflammation* | Lymphocytes | Macrophages | Neutrophils |  |  |  |  |  |  |  |  |  |  |  |  |
| P1 | 3 | <5 | 1.0 | 0.0 | 1.0 | 1.0 | 0.0 | 0.0 | 0.0 | 1.0 | 0.0 | 0.0 | 0.0 | 1.0 | 0.0 | 0.0 | 0.0 | 0.0 |
| P2 | 3 | <5 | 1.0 | 1.0 | 1.0 | 1.0 | 0.0 | 0.0 | 0.0 | 1.0 | 0.0 | 0.0 | 0.0 | 1.0 | 0.0 | 0.0 | 0.0 | 0.0 |
| P3 | 3 | 5 | 1.0 | 1.0 | 1.0 | 1.0 | 0.0 | 0.0 | 0.0 | 1.0 | 0.0 | 0.0 | 0.0 | 0.0 | 0.0 | 0.0 | 0.0 | 0.0 |
| T1 | 3 | 10 | 1.5 | 1.0 | 1.0 | 1.0 | 0.0 | 0.0 | 0.0 | 1.0 | 0.0 | 0.0 | 0.0 | 1.0 | 0.0 | 0.0 | 0.0 | 0.0 |
| T2 | 3 | 25 | 3.0 | 2.5 | 2.0 | 3.5 | 3.5 | 4.0 | 0.0 | 3.0 | 2.0 | 1.5 | 0.0 | 2.0 | 2.0 | 3.8 | 1.0 | 0.8 |
| T3 | 3 | 15 | 2.0 | 2.0 | 2.0 | 2.0 | 0.0 | 0.0 | 0.0 | 2.0 | 1.0 | 0.0 | 1.0 | 1.0 | 0.0 | 0.0 | 0.5 | 0.5 |
| C1 | 3 | 15 | 2.5 | 2.0 | 2.0 | 3.0 | 2.5 | 3.0 | 0.0 | 3.0 | 1.0 | 2.0 | 1.0 | 1.0 | 2.0 | 3.0 | 0.5 | 1.5 |
| C2 | 3 | 20 | 2.5 | 2.5 | 2.0 | 2.5 | 2.5 | 3.0 | 0.0 | 2.0 | 1.0 | 0.0 | 3.0 | 1.5 | 2.0 | 3.0 | 0.5 | 1.5 |
| C3 | 3 | 20 | 2.5 | 2.0 | 2.5 | 2.5 | 2.0 | 2.0 | 0.0 | 3.0 | 2.0 | 0.0 | 2.0 | 1.0 | 2.0 | 2.0 | 1.0 | 1.0 |
| P4 | 5 | 5-10 | 2.0 | 2.0 | 2.0 | 1.5 | 0.0 | 0.0 | 0.0 | 2.0 | 2.0 | 0.0 | 1.5 | 1.0 | 0.0 | 0.0 | 1.0 | 0.8 |
| P5 | 5 | <5 | 1.0 | 1.0 | 1.0 | 1.0 | 0.0 | 0.0 | 0.0 | 1.0 | 0.0 | 0.0 | 1.0 | 1.0 | 0.0 | 0.0 | 0.0 | 0.5 |
| P6 | 5 | 15 | 2.0 | 2.0 | 2.0 | 2.0 | 0.0 | 0.0 | 0.0 | 2.0 | 2.0 | 0.0 | 0.0 | 2.0 | 0.0 | 0.0 | 1.0 | 0.0 |
| T4 | 5 | 5-10 | 2.0 | 2.0 | 1.5 | 1.0 | 0.0 | 0.0 | 0.0 | 1.0 | 2.0 | 0.0 | 0.0 | 2.0 | 0.0 | 0.0 | 1.0 | 0.0 |
| T5 | 5 | 25 | 3.0 | 3.0 | 2.0 | 3.0 | 2.0 | 2.0 | 0.0 | 3.0 | 3.0 | 2.0 | 2.0 | 3.0 | 3.0 | 2.0 | 1.5 | 2.0 |
| T6 | 5 | 0 | 0.0 | 0.0 | 0.0 | 0.0 | 0.0 | 0.0 | 0.0 | 0.0 | 0.0 | 0.0 | 0.0 | 0.0 | 0.0 | 0.0 | 0.0 | 0.0 |
| C4 | 5 | 80 | 4.0 | 4.0 | 3.0 | 3.0 | 3.0 | 3.0 | 3.0 | 3.0 | 4.0 | 3.0 | 3.5 | 3.0 | 3.5 | 3.0 | 3.5 | 3.3 |
| C5 | 5 | 15-20 | 2.5 | 3.0 | 2.5 | 2.0 | 2.0 | 2.0 | 1.5 | 2.0 | 2.0 | 2.5 | 0.0 | 3.0 | 2.0 | 2.0 | 1.8 | 1.3 |
| C6 | 5 | 50 | 4.0 | 3.5 | 3.0 | 3.0 | 3.0 | 3.0 | 4.0 | 2.5 | 4.0 | 3.5 | 4.0 | 3.5 | 3.0 | 3.0 | 4.0 | 3.8 |
| P7 | 13 | 20 | 2.0 | 2.0 | 2.0 | 1.0 | 0.0 | 0.0 | 0.0 | 1.5 | 0.0 | 1.5 | 0.0 | 1.5 | 0.0 | 0.0 | 0.8 | 0.8 |
| P8 | 13 | 30 | 2.0 | 2.0 | 2.0 | 2.0 | 0.0 | 0.0 | 0.0 | 1.5 | 0.0 | 0.0 | 0.0 | 2.0 | 0.0 | 0.0 | 1.0 | 0.0 |
| P9 | 13 | 30 | 2.0 | 2.0 | 2.0 | 2.0 | 0.0 | 1.0 | 1.0 | 1.5 | 0.0 | 0.0 | 0.0 | 2.0 | 0.0 | 0.5 | 1.5 | 0.0 |
| T7 | 13 | 15 | 2.0 | 2.0 | 2.0 | 1.5 | 0.0 | 0.0 | 0.0 | 1.5 | 0.0 | 0.0 | 0.0 | 2.0 | 0.0 | 0.0 | 1.0 | 0.0 |
| T8 | 13 | 5 | 1.0 | 1.5 | 1.0 | 1.0 | 0.0 | 0.0 | 0.0 | 0.0 | 0.0 | 0.0 | 0.0 | 1.0 | 0.0 | 0.0 | 0.5 | 0.0 |
| T9 | 13 | 20 | 1.5 | 1.5 | 1.0 | 1.5 | 0.0 | 0.0 | 0.0 | 1.0 | 0.0 | 0.0 | 0.0 | 1.5 | 0.0 | 0.0 | 0.8 | 0.0 |
| C7 | 13 | 5 | 1.0 | 1.0 | 1.0 | 1.0 | 0.0 | 0.0 | 0.0 | 0.0 | 0.0 | 0.0 | 0.0 | 1.5 | 0.0 | 0.0 | 0.8 | 0.0 |
| C8 | 13 | 40 | 2.0 | 2.0 | 2.0 | 2.0 | 0.0 | 1.0 | 1.0 | 1.5 | 0.0 | 0.0 | 1.5 | 2.0 | 0.0 | 0.5 | 1.5 | 0.0 |
| C9 | 13 | 10 | 1.0 | 1.0 | 1.0 | 1.0 | 0.0 | 1.0 | 0.0 | 0.0 | 0.0 | 0.0 | 0.0 | 1.5 | 0.0 | 0.5 | 0.8 | 0.0 |

Histopathological scoring of formalin-fixed paraffin-embedded lung tissue from hamsters of experimental groups as abbreviated (P = prophylactic, T = therapeutic, C = control) at indicated days post infection (dpi). Degree of inflammation <sup>(\*)</sup> is scaled as (1) minimal, (2) mild,
(3) moderate or (4) severe. For all other parameters rating refers to occurrence rate from (1) sporadic, (2) mild, (3) moderate to (4) severe.
Scores were assessed as means from parameters with corresponding letters as indicated.

